# mBAT-combo: a more powerful test to detect gene-trait associations from GWAS data

**DOI:** 10.1101/2022.06.27.497850

**Authors:** Ang Li, Shouye Liu, Andrew Bakshi, Longda Jiang, Wenhan Chen, Zhili Zheng, Patrick F. Sullivan, Peter M. Visscher, Naomi R. Wray, Jian Yang, Jian Zeng

**Author notes:** Correspondence: Jian Zeng.

## Abstract

Gene-based association tests aggregate multiple SNP-trait associations into sets defined by gene boundaries. Since genes have a direct biological link to downstream function, gene-based test results are widely used in post-GWAS analysis. A common approach for gene-based tests is to combine SNPs associations by computing the sum of *χ*^2^ statistics. However, this strategy ignores the directions of SNP effects, which could result in a loss of power for SNPs with masking effects (e.g., when the product of the effects of two SNPs and their linkage disequilibrium (LD) correlation is negative). Here, we introduce “mBAT-combo”, a new set-based test that is better powered than other methods to detect multi-SNP associations in the context of masking effects. We validate the method through simulations and applications to real data. We find that of 35 blood and urine biomarker traits in the UK Biobank, 34 traits show evidence for masking effects in a total of 4,175 gene-trait pairs, indicating that masking effects in complex traits is common. We further validate the improved power of our method in height, body mass index and schizophrenia with different GWAS sample sizes and show that on average 95.7% of the genes detected only by mBAT-combo with smaller sample sizes can be identified by the single-SNP approach with larger sample sizes (average sample size increased by 1.7-fold). For instance, *LRRC4B* is significant only in our method for schizophrenia, which has been shown to play a role in presynaptic pathology using genetic fine-mapping and evidence-based synaptic annotations. As a more powerful gene-based method, mBAT-combo is expected to improve the downstream pathway analysis or tissue and cell-type enrichment analysis that takes genes identified from GWAS data as input to understand the biological mechanisms of the trait or disease. Despite our focus on genes in this study, the framework of mBAT-combo is general and can be applied to any set of SNPs to refine trait-association signals hidden in genomic regions with complex LD structures.

## Introduction

Genetic variants identified from genome-wide association studies (GWAS) of complex traits are often collapsed to genes (defined as the minimum start and maximum end boundaries over all identified gene transcripts) to seek biological interpretation of trait-variant associations and conduct post-GWAS analysis. One approach to generating an association statistic for a gene is to select an index SNP with the smallest P-value^1,2^. Alternatively, a gene-based test that aggregates association signals across all SNPs within the boundaries of a gene^3^ can be more powerful for the following reasons. First, it is common to observe multiple causal variants with small effects in a region^4–6^. Even if there is a single but unobserved causal variant, it may not be perfectly tagged by any individual common SNP, especially if it is uncommon or rare. In either case, aggregating signals over SNPs in the set can improve power. Second, compared to GWAS, gene-based association tests have orders of magnitude fewer significance tests compared to GWAS (approximately 20K versus 8 million) and so the threshold for declaring a significant test requires less correction for multiple testing. Genes identified by gene-based tests have been used to explain trait-variant associations^7^, put forward to downstream gene-set or pathway analysis^8,9^, and used to infer causative tissues and cell types for complex traits and diseases in conjunction with functional annotations^10,11^ or single-cell RNA-seq data^12,13^.

State-of-the-art methods for gene-based test only require GWAS summary statistics (i.e., SNP association z-scores or *χ*^2^ statistics), usually with a reference genotype dataset for computing the linkage disequilibrium (LD) correlation structure between SNPs (given extensive LD differences between ancestries^14^, it is important to match reference and GWAS populations in ancestry). A popular strategy to combine signals across common SNPs is to compute the sum of the *χ*^2^ statistics (referred to as the sum-*χ*^2^ strategy hereafter). VEGAS^15^, fastBAT^16^, and MAGMA^17^ are widely used methods that employ this strategy, with the main difference between them in how P-values are computed. The sum-*χ*^2^ strategy can lead to improved power, especially for genes harbouring multiple causal variants^16^. However, since this strategy is based on the sum of *χ*^2^ variables of individual SNPs, the method loses power when the underlying causal variants mask the effects of each other because of the correlation structure between them. In single SNP association tests this issue can be addressed through the conditional and joint (COJO) analysis algorithm^18^.

A standard GWAS reports the per-SNP *χ*^2^ which is a measure of the marginal association for the trait, regardless of all other SNPs. If there are multiple causal variants at a locus, the power of detecting marginal associations can substantially diminish if the product of the causal effects and the LD correlation between causal variants is negative (i.e., negative genetic covariance among causal variants)^19^. This phenomenon is referred to as “masking”, that is, the masking of causal effects in marginal analysis due to LD. LD is often parameterised using *r*^2^, but here we use the directional correlation (*r*). Fig.1 illustrates the masking effect concept considering two variants in positive LD with opposite effects, where the marginal effect of either variant, estimated from a single-SNP model, is smaller than the true effect because the two causal effects tend to be cancelled out with each other by LD. Similarly, the true effects can be masked when the two causal variants have the same effect direction but negative LD. Given the mounting evidence for the presence of multiple causal variants at a locus^4–6^ and the action of long-term stabilising selection in human traits^20,21^ (or negative selection on trait-associated variants)^22–25^, negative LD could be created between trait-increasing alleles (see below for more discussion). We seek to test if masking effects are prevalent across the human genome.

**Figure 1.**
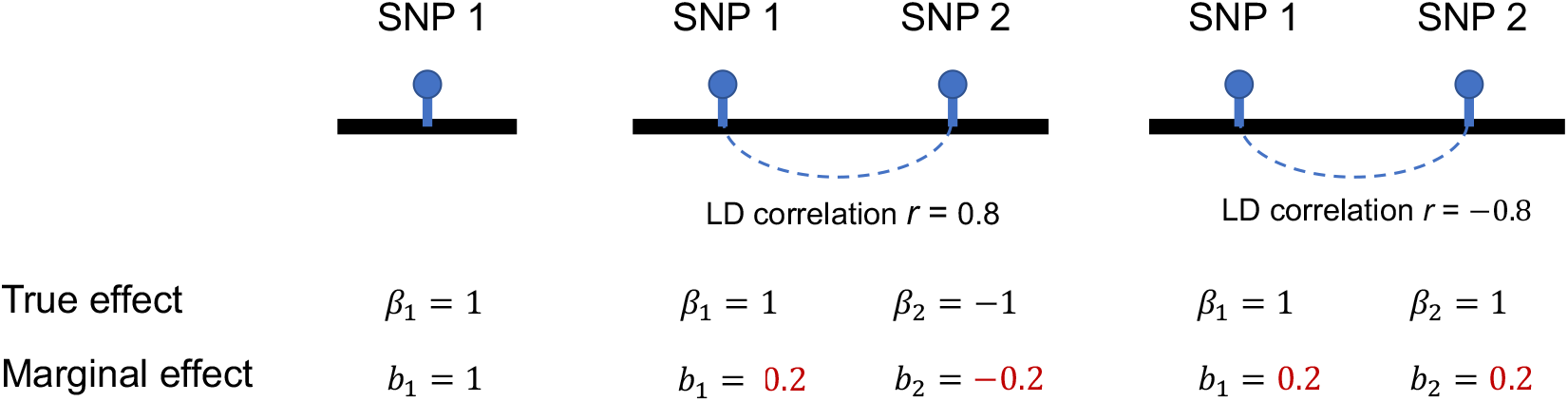
Schematic of masking effect. The masking effect occurs when the product of the true effects and the LD correlation is negative. The marginal effect of each SNP can be written as the sum of true effects weighted by the LD correlation between the two SNPs: b_i_ = β_i_ + r × β_j_. The marginal effect size is smaller than the true effect size when the SNP is masked by another SNP in LD with it.

While it is difficult to disentangle genetically correlated variants in the marginal analysis, joint analysis based on a multiple regression approach can help uncover signals of masked variants. There are two general types of joint analyses. One type of analysis performs the hypothesis test on the joint associations of multiple variants on the trait. Brown et al.^19^ proposed a pairwise testing between every pair of SNPs in a sliding window to detect masking effects. Yang et al.^4^ employed a more sophisticated algorithm to include variants in the joint-association test by a stepwise model selection, which searches for secondary signals that are conditionally independent of the most genome-wide significant SNP in a region (i.e., COJO). However, SNP hypothesis testing requires a stringent significant threshold to correct for genome-wide multiple testing correction, and it is not devised for set-based tests (e.g., for genes). Another type of joint analysis is based on estimation of genetic variance. While the total genetic variance is typically estimated by a mixed model approach, it is possible to estimate the local genetic variance with a fixed effect model that simultaneously fit all SNPs in the focal region^26^. The variance explained by all SNPs would capture the masking effect because it accounts for the LD between SNPs. Therefore, the estimate can be greater than the sum of that estimated for each independently genome-wide significant SNP^27^, contributing to filling the gap of “missing heritability”^28–30^. Although the variance estimation approach is expected to be more powerful than the SNP association test, this approach was proposed for applications on mega-base sized genomic segments (the median size of protein-coding genes is 27.4kb and only 0.3% are > 1 Mb); estimates would thus tend to be biased when applying to gene regions^26^.

In this study, we proposed a novel gene-based method, <u>m</u>ultivariate set-<u>B</u>ased <u>A</u>ssociation <u>T</u>est (mBAT), to improve power of identifying genes harbouring variants with masking effects. Our method is derived based on the quadratic form theory in normal variables^31^, and only requires GWAS summary statistics and an appropriate LD reference. In essence, mBAT tests for the non-zero variance explained by the SNPs in the set, thus it has the advantage of high power as featured in the variance estimation approach. Meanwhile, our test statistic is analytically tractable, with closed-form solutions for the P-values. We show by extensive simulations that mBAT has greater power than the sum-*χ*^2^ strategy (taking fastBAT and MAGMA as examples) in the presence of masking effects. To maximise overall power regardless of the masking effects, we further proposed a hybrid method, mBAT-combo, by combining mBAT and fastBAT test statistics through a Cauchy combination method, a recently developed method to combine different test statistics without knowing the correlation structure. We applied mBAT and mBAT-combo to GWAS results for a range of complex traits of different genetic architecture^32^ to compare the performance of COJO, MAGMA and fastBAT. For height, body mass index (BMI), and schizophrenia (SCZ), we investigated improved power empirically by comparing results from GWAS with smaller sample sizes. Our methods have been implemented in the GCTA software ^33^ and in a R package (https://github.com/Share-AL-work/mBAT).

## Methods

### mBAT

Let **x** be a *m* × 1 vector of random variables that follow a multivariate normal distribution with mean zero and variance-covariance **∑, x**∼*N*(**0, ∑**). Suppose **A** is a *m* × *m* symmetric matrix. Then, the quadratic form of **x** in association with matrix **A** is^34^

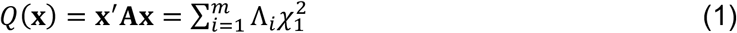

where Λ_*i*_ is the *i*^th^ eigenvalue of **∑A** in descending order, and 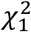 is an independent central chi-squared variable with 1 degree of freedom (d.f.). If **A = ∑**^−1^, then **∑A = I**, and thus Λ_*i*_ = 1. It follows that

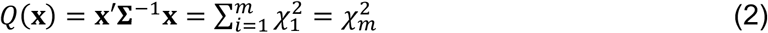

where 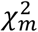 is a central chi-squared variable with d.f. = *m*. In the context of GWAS, under the null hypothesis that none of the *m* SNPs in the set is associated with the trait, the z-scores of SNPs would have the same distribution as **x**, with **∑** being the LD correlation matrix among SNPs (denoted as **R)**:

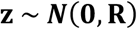

The test statistic of mBAT is

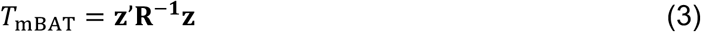

which, according to Eq (2), has a central chi-squared distribution:

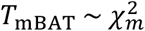

See the **Supplementary Note** for a direct proof that the mBAT test statistic follows a central chi-squared distribution. Note that because the normality assumption of **z** always holds under the null, mBAT applies to genes with different genetic architectures.

We show below that essentially mBAT tests if the variance explained by all SNPs in the set is zero. Let **X** be the *n* × *m* standardised genotype matrix with column mean zero and variance one (*n* is the GWAS sample size), and ***β*** be the *m* × 1 vector of the true effects of SNPs. The variance explained by the SNPs is

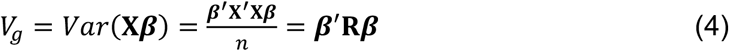

Let **b** be the vector of SNP marginal effect estimates from GWAS. We have

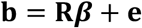

where 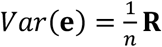 assuming unit phenotypic variance and each individual *b*_*j*_ explaining a negligible proportion of variance. It can be seen that **b** converges to **R*β*** as *n* increases to infinity, which gives

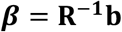

Substitution in Eq (4) gives

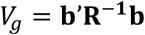

Given that the sampling variance of *b*_*j*_ is 1/*n* as shown above, the mBAT test statistic is simply a product of the genetic variance and the GWAS sample size:

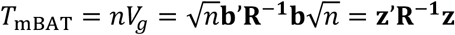

Thus, mBAT can be regarded as a test for a nonzero variance explained by the SNPs in the gene region.

One challenge in the computation of the mBAT test statistic is that the **R**^−1^ may not always exist in a gene region because the LD matrix for a set of consecutive SNPs is often rank deficient. Thus, we used the pseudo-inverse of the **R, R**^−^ **= UD**^−1^**U**′, where **D** = *diag*{*λ*_*i*_} with *i* = 1, …, *p* is a diagonal matrix of the *p* positive eigenvalues of **R**, and **U** is a *m* × *p* orthogonal matrix whose columns are eigenvectors corresponding to the eigenvalues in **D**. As a result, the test statistic follows a chi-squared distribution with d.f. equal to the number of selected eigenvalues (**Supplementary Note**):

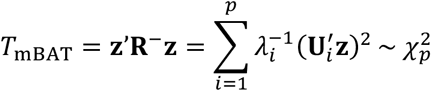

In practice, **R** would be computed from a reference sample if individual-level genotype data from GWAS are not available. To remove LD differences between the reference and GWAS samples due to sampling, for each gene region, we chose to only include the top *k* principal components needed to explain at least *γ*% of the total variance in the LD matrix, where the total variance of the LD matrix is computed as the sum of all positive eigenvalues. Depending on the LD structure, *k* can differ between genes. We investigated different thresholds for proportion of variance explained *γ* = 0.7, 0.75, 0.8, 0.85, 0.9, 0.95, 0.99 and 1 in simulations, and determined to use 0.9 in real data analyses.

### The sum-*χ*^2^ strategy

The connection between the sum-*χ*^2^ strategy and mBAT can be regarded as the use of different **A** matrices in Eq (1). If we set **A = I**, the *m* × *m* identity matrix, the quadratic form of z-scores is

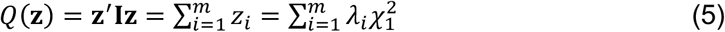

Thus, both Eq (2) and (5) can be regarded as a special case of the general form of Eq (1). This is the test statistic used in fastBAT, VEGAS and MAGMA. Unlike mBAT, it does not have a closed-form solution for computing the P-value, i.e., Pr(*Q*(**z**) > *q*) with *q* being the observed value. fastBAT uses the Saddlepoint method for an approximation^16,35^. VEGAS calculates the P-value by simulation which is computational demanding^15^. MAGMA uses Imhof’s method to compute the P-value by numerical integration but when it fails, an empirical P-value from simulations is reported instead^17,36^ (in which case the minimum P-value is 1/number of simulations).

### mBAT-combo

The relative superiority of mBAT or the sum-*χ*^2^ strategy could depend on the genetic architecture of the gene for the trait. To achieve the optimal power, we combine the test results of mBAT and fastBAT by the aggregated Cauchy combination method (ACAT), an established omnibus test^37^ that combines P-values from different tests without knowing the correlation structure among the test statistics. Let *P*_*i*_ be the P-value from fastBAT or mBAT. With equal weights assigned to the two tests, we have

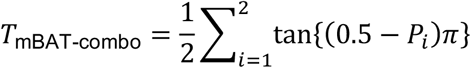

Where tan{(0.5 – *P*_*i*_)π} follows a standard Cauchy distribution^37^ given that *P*_*i*_ is uniformly distributed under the null^38^. The P-value of mBAT-combo is expected to approach the smallest P-value from mBAT or fastBAT in the presence of a true signal, therefore giving the highest power overall. When one of the methods give an extremely small P-value (i.e., < 1×10^−16^), we have

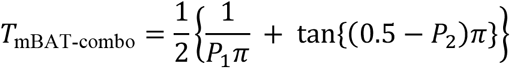

Where *P*_1_ and *P*_2_ denote the P-values smaller and larger than the threshold, respectively. When both methods’ P-values are smaller than the threshold, we have

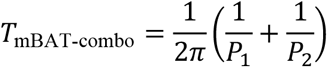

### Derivation of theoretical power

To compare analytically the performance of mBAT and methods using the sum-*χ*^2^ strategy, we derived the theoretical power given a false positive rate for the two approaches. It requires, for each method, knowledge of the distribution under the null model and evaluating Pr(*T* > *τ*) under the alternative model, where *T* is the test statistic and *τ* is the quantile value given a false positive rate *α* under the null. Without loss of generality, let us consider a simple case with two (*m* = 2) causal variants in LD with each other. The causal effects are *β*_1_ and *β*_2_, respectively, and the LD between the two variants is *r*.

The mBAT test statistic follows a central chi-squared distribution with d.f. = 2 under the null and a non-central chi-squared distribution with d.f. = 2 under the alternative, where the non-centrality parameter 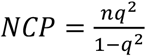 with 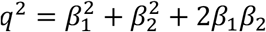 being the variance explained by the two causal variants jointly.

The sum-*χ*^2^ test statistic does not have a known distribution either under the null or the alternative model. However, it has been shown that the distribution of the sum of correlated central chi-squared variables can be well approximated by a Gamma distribution^39^.

Specifically,

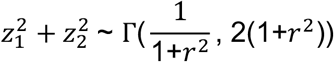

where *r*^2^ is the squared correlation between the two chi-squared variables. The quantile value *τ* given *α* can be computed by the inverse of the Gamma cumulative density function (CDF). Under the alternative model, the test statistic is the sum of two correlated non-central chi-squared variables with d.f. = 1, 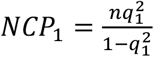 and 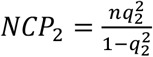, respectively, where 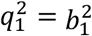 with *b*_1_ = *β*_1_ *+ rβ*_2_ and 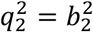 with *b*_2_ = *β*_2_ *+ rβ*_1_. It is known ^39^ that this statistic can be transformed into the sum of two independent non-central chi-squared variable with d.f. = 1 and NCP equal to 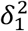 and 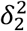, respectively, where

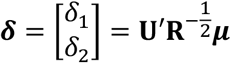

and 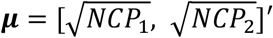. Then, the Saddlepoint approximation^39^ can be used to compute the power, i.e., Pr(*T* > *τ*), without knowing the exact distribution under the alternative. An R script for computing the theoretical power in the case of two causal variants can be found at GitHub (https://github.com/Share-AL-work/mBAT).

### Proof-of-concept simulations

We used simulations based on simulated genotypes to contrast the performance of our methods to the sum-*χ*^2^ strategy (fastBAT was used as a representative) using a minimal number of parameters to depict the scenarios of masking and non-masking effects. We considered two SNPs with simulated genotypes from a bivariate normal distribution, 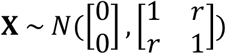, where the correlation between the genotypes (*r*, i.e., LD) varied from 0.1 to 0.99. The effect of SNP 1 (*β*_1_) was set to 1. The effect of SNP 2 (*β*_2_) varied from −0.1 to −1. This setting represented a scenario of masking effects, i.e., *β*_1_ × *β*_2_ × *r* < 0. In the scenario of non-masking effects, the direction of *β*_2_ was set to be positive. We simulated genotypes for 500 individuals, each had a genotypic value *g*_*i*_ = *X*_*i*1_*β*_1_ *+ X*_*i*2_*β*_2_ and a phenotypic value *y*_*i*_ = *g*_*i*_ *+ e*_*i*_, with *e*_*i*_ ∼ *N*(0, 100). The marginal effect of each SNP was estimated by regressing the phenotypes on the SNP genotypes. The z-scores of marginal associations and the true LD correlation matrix were the input data for each method. This simulation process was repeated 1,000 times for each combination of *β*_2_ and *r* values. An R script for running the proof-of-concept simulations can be found at GitHub (https://github.com/Share-AL-work/mBAT).

### Simulations based on real genotypes

We performed simulations based on real genotypes to 1) determine the optimal *γ* parameter (the proportion of variance in LD explained by the chosen eigenvalues) for subsequent real data analyses, and 2) compare our methods with fastBAT and MAGMA in terms of false positive rate (FPR; the proportion of null genes with P-values < 0.05) and power (the proportion of causal genes identified with significant P-values). We defined a gene unit as a SNP set located in ±50 Kb of base-pair positions for the start and stop codons for each gene derived from gene ID and annotation list released from GENCODE in the genome build of hg19/GRch37 Release 37 (GRCh37) (https://www.gencodegenes.org/human/). The simulations were conducted based on 100 randomly selected protein-coding genes on chromosome 1 with 45,657 imputed common SNPs (MAF > 0.01) on a random sample of 34,993 unrelated individuals of European ancestry in the UKB. All the selected genes have at least 50 SNPs to ensure that we had enough room to sample up to 8 causal variants for each gene. Under the null hypothesis, individual phenotypes were sampled from a standard normal distribution without a genetic component. Under the alternative hypothesis, 100 genes were sampled as the causal genes in the scenarios of either masking or non-masking effects (the same definition as in the proof-of-concept simulation above), with a trait heritability *h*^2^ = 0.1. We extracted 2-8 causal variants per gene from SNPs that were located in the region unique to the gene. We further simulated different levels of statistical power (measured by NCP=50, 100 or 150) by performing simulation for each gene in isolation with 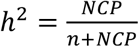. The simulation was repeated 1,000 times under the null hypothesis and 10 times for each of the settings under the alternative hypothesis. In the analysis, the LD correlation matrix was computed from either the GWAS sample (internal LD) or an independent dataset (external LD). The unrelated individuals of European ancestry in the Genetic Epidemiology Research on Aging (GERA) dataset^40^ (n= 51,258) or 1000 Genome Project dataset^41^ (n= 494) were used as the external LD data sets to investigate the influence of reference sample size on our methods. Finally, we randomly flipped the direction of the marginal effect for 0.5%, 1%, 2%, 5%, or 7% of the SNPs to examine the robustness of our methods to the potential allelic errors in the GWAS summary data.

### Real data analyses

First, we benchmarked different methods with multiple linear regression (MLR) using individual-level genotype and phenotype data in UKB height. MLR was regarded as the gold standard because it invoked an ANOVA test for the variance explained by all SNPs in the set, which accounts for the masking effects, but can only be applied when individual-level data are available. The linear regression model (LRM) implemented in fastGWA^42^ was used to conduct GWAS analyses on 8,531,416 variants with MAF ≥ 0.01 in all the UKB unrelated individuals of European ancestry with height phenotypes (n = 348,423). Age, sex, and the top 20 PCs provided by the UKB were used as covariates in fastGWA-LRM and MLR. In all the analyses of real data, we always ran mBAT, mBAT-combo, fastBAT v1.93.2 and MAGMA v1.09 using the imputed genotype data from GERA as the LD reference. For this benchmark analysis, we also ran these methods using the internal LD as a sensitivity check. MLR was carried out using the top *k* eigenvectors obtained from the singular value decomposition (SVD) of the genotype matrix for each gene as independent variables, with *k* being the same number of eigenvalues corresponding to the *γ* parameter in mBAT (*γ* was chosen to be 0.9 for the real data analyses).

Next, we applied COJO analysis to the GWAS summary data from 35 UKB blood and urine biomarker traits using genotype data from all UKB unrelated individuals as LD reference. We used COJO to identify genes with significant masking effects and compared the power of detecting such putative masking genes between different gene-based methods. More specifically, a gene is identified as a putative masking gene if it had at least 2 joint SNP effects detected by COJO, with the top associated SNP at the genome-wide significant level of *P*_joint_ < 5×10^−8^, a secondary SNP at a relaxed significant level of *P*_joint_ < 5×10^−6^, and the genetic covariance between the two SNPs being negative (i.e., *β*_1_ × *r β*_2_ < 0). While we knew COJO was underpowered to identify all masking genes, those identified were unlikely to be false given the stringent significance threshold imposed by multiple test correction at SNP level.

Finally, we applied mBAT and mBAT-combo to the publicly available summary data from the latest and earlier releases of GWAS meta-analyses for height^5,43^, BMI^5,44^ and SCZ^45,46^ with different sample sizes. A gene identified only by mBAT or mBAT-combo was called if it passed genome-wide significance threshold at gene level (*P*_mBAT_ < 2.7×10^−6^) and there was no genome-wide significant SNP (*P*_GWAS_ > 5×10^−8^) within 1MB of the gene. We further investigated the results that were only significant in mBAT (mBAT-combo) but neither in fastBAT nor in MAGMA. We used the results from analyses applied on height, BMI and SCZ and sought to find whether most of the genes identified only by mBAT (or mBAT-combo) in smaller samples could be discovered by GWAS with larger sample sizes (presence of a genome-wide significant SNP in gene region ± 50kb).

## Results

### Method comparison by proof-of-concept simulations

The proof-of-concept simulations found no inflation in FPR for mBAT, fastBAT (as a representative sum-*χ*^2^ strategy like VEGAS and MAGMA) or mBAT-combo (**Fig. 2a**). We derived the theoretical power for mBAT and fastBAT (**Fig. 2c** and **2d**, solid line; **Methods**), and the results were consistent with those computed empirically across simulation replicates (**Fig. 2c** and **2d**, dashed line). These results confirmed that mBAT was superior to fastBAT in the presence of masking effects, especially when the two causal variants had similar magnitudes of effect sizes. The benefits of mBAT increased with LD up to the point where LD was so high that the two variants effectively collapsed into one (**Fig. 2c**). When the effect sizes of the two variants became distinctive or the variants were in lower LD, the difference between mBAT and fastBAT diminished because of a smaller masking effect between the two variants, i.e., a decreased proportion of genetic variance being masked (**Fig. S1**). We noted that fastBAT gave a slightly higher power than mBAT under the scenario of non-masking effect **(Fig. 2d**). This is likely because the marginal association signals were enhanced when the genetic covariance is positive, which led to a stronger signal in the sum-*χ*^2^ strategy. Nevertheless, mBAT-combo, which combines the merits of both tests, gave a power as high as that of the best method in either scenario (**Fig. 2b**), with consistent performance across different settings (**Fig. 2c** and **2d**). As expected, the P-values from mBAT were highly consistent with those from MLR (**Fig. S2 and S3**), which were both more significant than fastBAT in the scenario of masking effect (**Fig. S2**).

**Figure 2:**
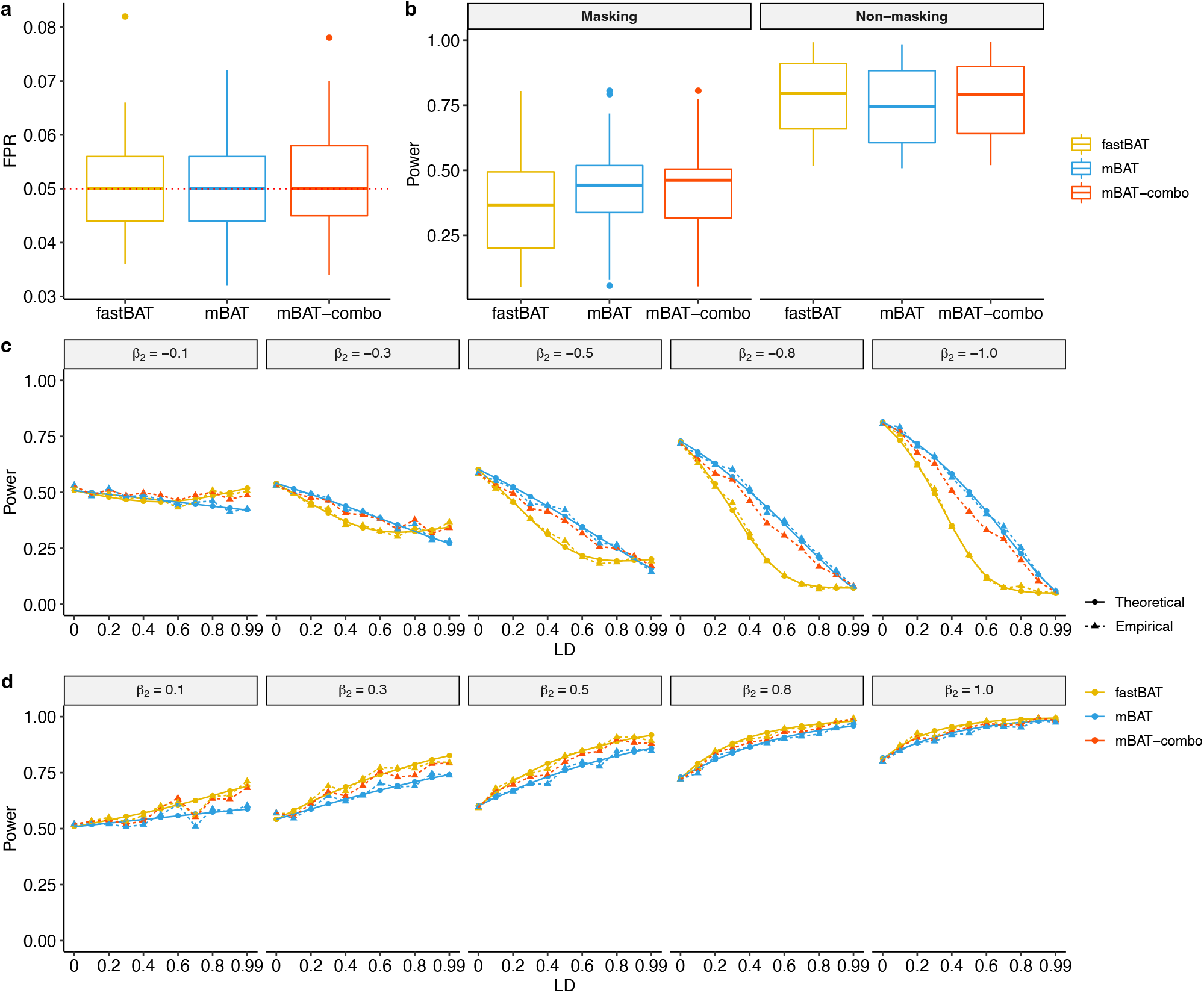
False positive rate (FPR) and power comparisons of mBAT, fastBAT and mBAT-combo using proof-of-concept simulations. a) FPR under the null model; b) Overall power under the causal model in scenarios of masking and non-masking effects. Each boxplot consisted of results across different settings for effect size of the second SNP (effect of the first SNP was set to be 1) and the LD correlation between the two SNPs, with 1,000 simulation replicates per setting. c) and d) Theoretical power (solid line) and empirical power (dashed line; mean over 1,000 replicates) given a P-value of 0.05 in the scenario of masking effect (c) and non-masking effect (d) with different settings for the second SNP effect size (grey header boxes) and LD levels (x-axes). Results of different methods are shown by colours.

### Assessing performance in simulations based on real genotypes

We calibrated the critical parameter in mBAT, the proportion (*γ*) of variance in the LD matrix explained collectively by the selected eigenvalues, by simulations based on genotype data in the UKB. Using an external LD reference from the GERA dataset, FPR was remarkedly inflated when *γ* = 1 (including all positive eigenvalues) and the inflation decreased when a smaller *γ* value was used (**Fig. S4a**). Thus, it is necessary to impose a filter on small eigenvalues which are trivial in variance explained but sensitive to sampling variation in LD between GWAS and reference samples as well as genotyping or allelic errors in GWAS. This is supported by our observations that mild LD pruning (pairwise r^2^ > 0.9) alleviated the inflation at *γ* = 1 (because complete LD may not persist across samples). Moreover, applying DENTIST^47^ (a quality control, QC, tool to detect LD heterogeneity in GWAS summary statistics) prior to the analysis could further reduce the inflation (**Fig. S4a**). Note that when *γ* is 0.95 or 0.90, FPR in mBAT had already been well-controlled regardless of the additional QC steps, and the removal of small eigenvalues also improved the robustness of mBAT in the presence of mislabelled alleles in GWAS summary statistics (**Fig. S5**). However, the highest power was achieved at *γ* = 0.9∼0.85 for mBAT and additional QC such as LD pruning or application of DENTIST reduced power (**Fig. S4b, S6 and S7**). Therefore, we chose *γ* = 0.9 for mBAT or mBAT-combo in the following analyses unless otherwise specified and without LD-pruning or application of DENTIST.

Next, we compared methods. By default, fastBAT performs pairwise LD-pruning at *r*^2^ > 0.9 and MAGMA implements a pruning of eigenvalues that cumulatively explain ≤ 0.1% of the variance in the LD matrix (equivalent to setting *γ* = 0.999). All methods were well calibrated under the null simulation. Under the causal model, mBAT had higher power than fastBAT under the masking scenario but not under the non-masking scenario (**Fig. 3**), consistent with the result from the proof-of-concept simulation. Overall, mBAT-combo test gave the highest power regardless of the masking or non-masking scenario, whereas MAGMA gave the lowest power compared to other methods across different settings (investigated in detail below). Similar patterns of performance were observed with different numbers of causal variants (*n*_QTL_ = 2, 4, 6 or 8), different levels of power (quantified by NCP per gene = 50, 100 or 150), and the use of internal LD or two external LD datasets with very different sample sizes (GERA (n= 51,258) and 1KG (n= 494) European samples datasets) (**Fig. S8-S10**).

**Figure 3.**
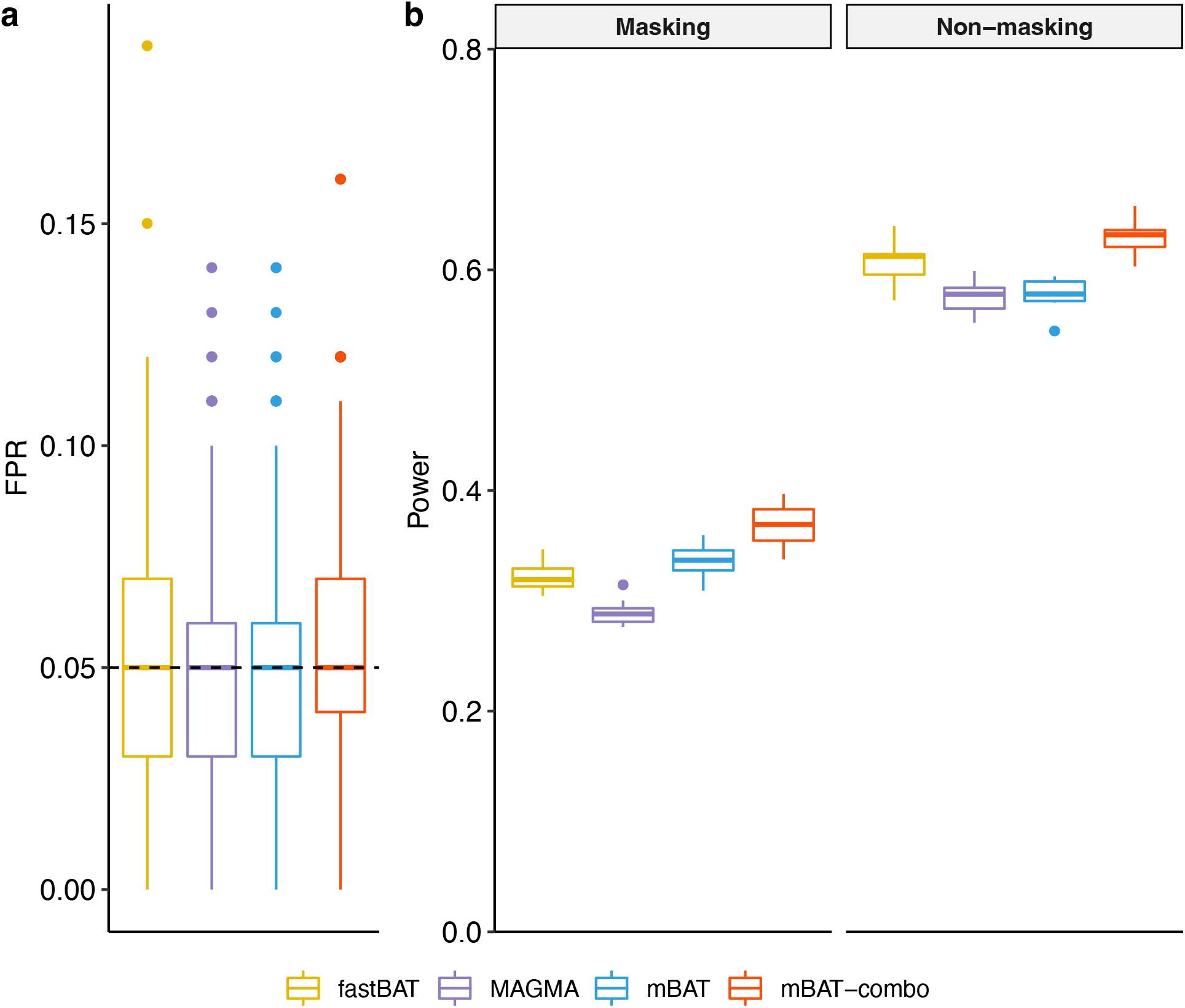
False positive rate (FPR) and power comparisons of different methods in simulations based on real genotype data. Colours represent different gene-based tests. a) FPR under the null model; Each boxplot represents the distribution of estimates across 1,000 simulation replicates; b) Overall power under the causal model in scenarios of masking and non-masking effects. Each boxplot represents the distribution of estimates across 10 simulation replicates.

### Benchmark with multiple linear regression in UKB height

In theory, mBAT tests for a nonzero genetic variance explained by all SNPs in the gene (**Methods**), and we have shown its equivalence to the multiple linear regression (MLR) approach by simple proof-of-concept simulations (**Fig. S2** and **S3**). Here, we investigate in the real data analyses the consistency between mBAT using GWAS summary statistics and MLR using individual-level genotype and phenotype data in the UKB height, in comparison with fastBAT and MAGMA (**Fig. 4**). Consistent with the simulation results, mBAT (or mBAT-combo) was almost identical to MLR when in-sample LD were used, and in strong agreement with MLR when using external LD (Pearson correlation *r* = 0.98), suggesting mBAT (or mBAT-combo) is robust to the choice of LD reference in practice once very small eigenvalues are eliminated (**Fig. S11**). Compared to the gold-standard MLR approach, both fastBAT and MAGMA tended to have compromised P-values, especially for genes with strong association signals, likely because of the presence of masking effects in those genes. In MAGMA when the Imhof integration method fails, P-values are generated through simulation, which limits the minimum P-value to 1/10^10^, where 10^10^ is the number of simulation replicates, and the Imhof’s method relies on numerical integration, which does not track the extreme P-values. If genes with extreme P-values (P < 5×10^−20^ in MLR) are excluded, then using external LD, mBAT (*r* = 0.98, regression slope *b* = 1.04) and mBAT-combo (*r* = 0.97, *b* = 1.03) remained most consistent with MLR, followed by fastBAT (*r* = 0.87, *b* = 0.72), and MAGMA had the least concordance (*r* = 0.79, *b* = 0.52) (**Fig. S12**).

**Figure 4.**
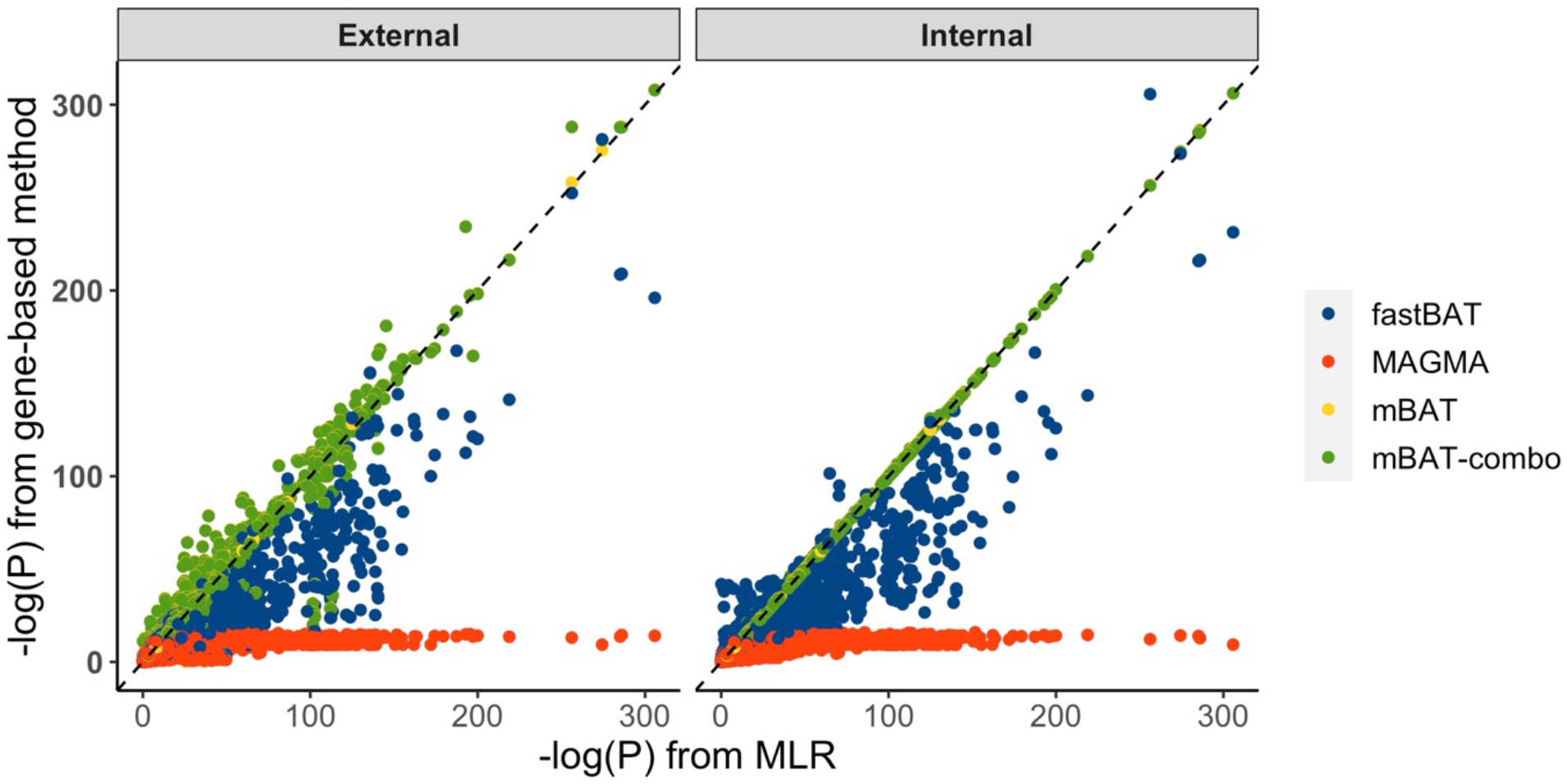
Benchmark of different gene-based test methods with multiple linear regression (MLR) in the UKB height, using external LD and in-sample LD. The y-axis is the -log10 P-values of different gene-based methods denoted by colours. The x-axis is the -log10 P-values of MLR with the same number of eigenvectors as that used in mBAT and mBAT-combo. See Supplementary Fig. 12 for a zoom-in when P_MLR_ > 5×10^−20^.

### Masking is common if not ubiquitous

Our hypothesis is that masking phenomenon is common, given the evidence of multiple causal variants in a locus and widespread negative selection in the human genome^4–6^. mBAT could uncover trait-associated genes that were missed by other methods due to masking effects, and so did mBAT-combo. We have shown that mBAT had relatively high power under masking, tested by simulations. To validate it in real data, we used the results from the COJO analysis (a conservative approach) to form a set of putative genes with masking effects and sought for two lines of evidence of validation. We applied our methods to 35 UKB blood and urine metabolite traits with diverse genetic architectures. mBAT-combo indeed identified more significantly associated genes than the sum-*χ*^2^ methods (**Fig. 5a**). On average across traits, mBAT-combo (mBAT) identified 19.7% (11.5%) more genes than fastBAT and 56.9% (43.9%) more than MAGMA.

**Figure 5.**
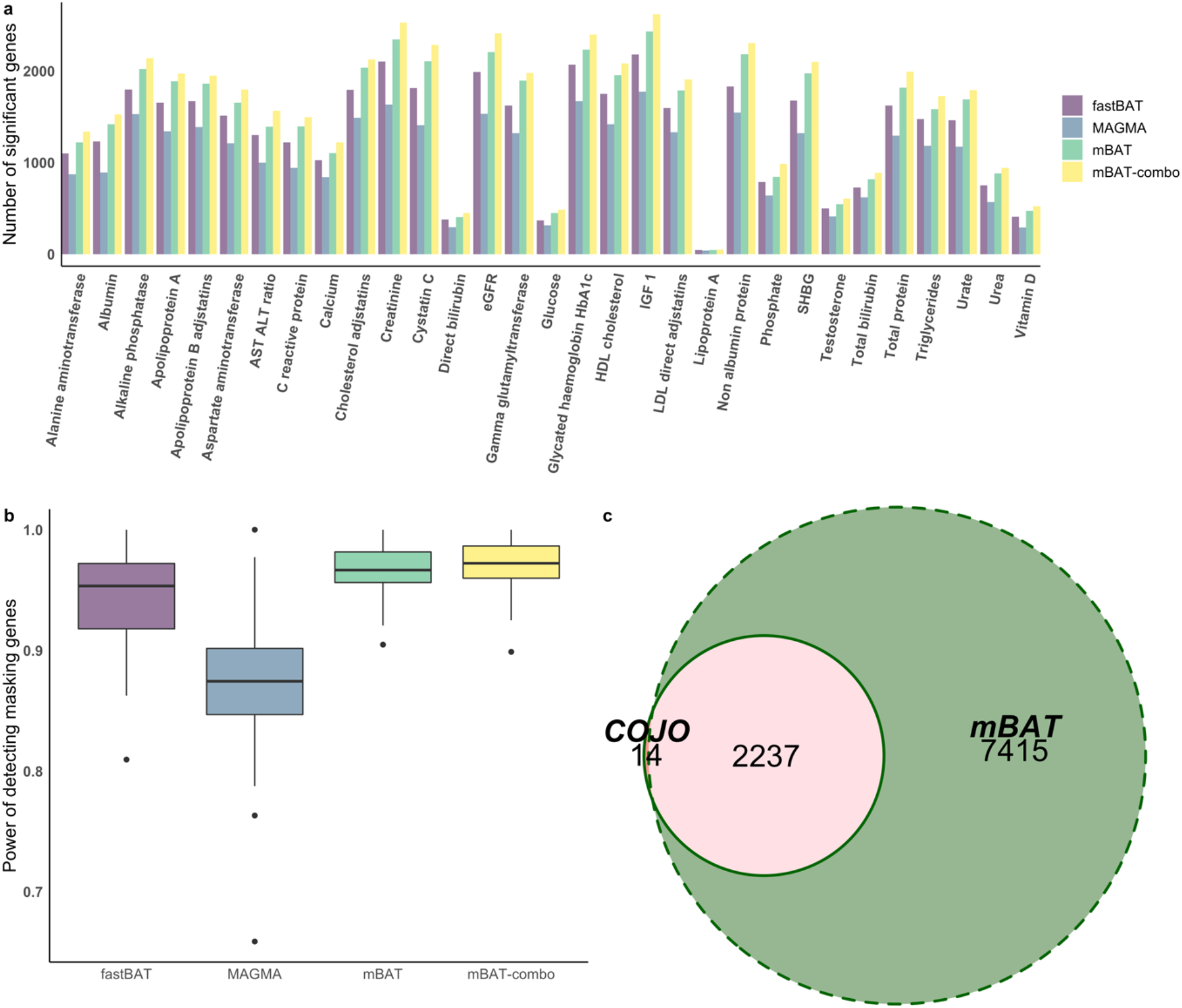
Improved power for identifying genes with masking effects for 35 blood and urine metabolite traits in the UKB. a) The total number of genes detected by different methods. b) Power of detecting putative genes with masking effects identified by COJO. Each boxplot represents the distribution of power estimates for each of the methods across the 35 traits. c) The numbers of genes identified by COJO with masking association signals, mBAT and the intersection.

First, we collected genes with evidence of masking effects from COJO, where the top two SNPs with significant joint-association signals had a negative product of their effects and LD correlation (**Methods**) and examined if mBAT had a higher power to detect these putative true positives. We found such putative genes with masking effects in 31/35 metabolite traits and the numbers varied from 21 to 402 across traits (**Fig. S13**), reflecting differences in trait genetic architecture. Of these 31 traits, mBAT-combo and mBAT had higher power detecting masking genes than other methods in 26 traits, leading to a significant improvement overall (t-test P-value = 3.4×10^−3^ between mBAT-combo and fastBAT and 2.9×10^−9^ between mBAT-combo and MAGMA) (**Fig 5b**). For genes with evidence of multiple causal variants but non-masking effects (the top-2 SNPs with significant COJO signals had a positive product of their effects and LD correlation), mBAT and mBAT-combo had a mean power as high as fastBAT but still significantly higher than MAGMA (**Fig. S14**). There were genes with multiple conditionally independent signals detected by COJO, except for creatinine, microalbumin, potassium and sodium in urine. In all methods, the power increased with an increasing number of COJO signals (**Fig. S15**), with mBAT and mBAT-combo always being the most powerful methods, consistent with the simulation result (**Fig. S9**).

The second line of evidence supporting the relatively high power of mBAT in detecting genes with masking effects is to investigate if genes that are only detected by mBAT (not significant in the sum-*χ*^2^ methods) are enriched with putative genes with masking effects (based on the COJO results). Across 34 traits, we found 4,273 gene-trait pairs that are conservatively significant in mBAT (at a more stringent threshold of P_mBAT_<5×10^−8^) but not significant in other methods after a standard Bonferroni correction within each trait (both P_fastBAT_ and P_MAGMA_ < 2.7×10^−6^). Among these gene-trait pairs, 115 (2.7%) harboured at least two joint association signals detected by COJO, of which the majority (98/115=85.2%) were consistent with the scenario of masking effects (**Table S1**). Namely, the COJO signals with masking effects were significantly enriched in genes that were only significant in mBAT (Fisher’s exact test P-value = 7.6×10^−20^, **Table S2**). Thus, association signals that are specifically detected by mBAT can be regarded as an evidence of masking effect associations.

Our results showed that of 35 blood and urine biomarker traits analysed, 34 had evidence for genes with masking effect associations (24 confirmed by COJO). Notably, mBAT had identified almost all putative masking genes defined by COJO (99.4%) but reported much more significant genes (**Fig. 5c**), suggesting a higher power of mBAT than COJO in terms of identifying masking genetic effects, in line with the simulation result (**Fig. S16**). All these results suggest that masking is common in trait-associated gene regions, and mBAT can improve power of detecting genes with masking effect in traits with diverse genetic architecture, thus making mBAT-combo the best method achieving an optimal power overall.

### Validation of novel genes found by mBAT-combo in GWAS data with larger sample sizes

We have shown by real trait analysis that mBAT (or mBAT-combo) has higher power in identifying genes with COJO signals of masking effects and such COJO signals were enriched in genes only detected by mBAT. Another approach to validate the novel genes identified by mBAT and mBAT-combo is to check if these genes go on to have a genome-wide significant SNP when the GWAS sample size increases. We performed this validation analysis in three complex traits (height, BMI and SCZ) for which GWAS summary statistics with different sample sizes are available. With mBAT-combo (mBAT), we identified 2,198 (2,111) novel genes for height (N = 241,427), 117 (104) for BMI (N = 226,289), and 576 (477) for SCZ (N_effective_= 57,231) at the genome-wide significance level (i.e., P < 2.7×10^−6^) using an earlier GWAS summary dataset for each trait^1,45^. A mBAT-combo (or mBAT) specific gene was claimed if it was significant in mBAT-combo (or mBAT) gene-based test, but no genome-wide significant SNP was found in the gene region ± 1Mb. Of these mBAT-combo (mBAT) specific genes, the vast majority can be validated in the subsequent GWAS with larger sample sizes (at least one SNP in the gene region ± 50kb reached genome-wide significance level of P_GWAS_ < 5×10^−8^), i.e., 99.3% (99.6%) for height (N= 456,426), 100% (99.1%) for BMI (N=456,426), and 87.2% (90.2%) for SCZ (N_effective_= 73,189) (**Fig. 6a-c**), with a pooled validation rate 95.5%(96.3%) across the three traits (by 1.9, 2.0 and 1.3 fold sample size increase respectively). For the mBAT-combo (mBAT) specific genes that could not be detected by either fastBAT or MAGMA, we also observed high validation rates, 99.0% (98.8%) for height, 100% (97.6%) for BMI, and 80.7% (76.7%) for SCZ (**Fig. 6d-f**). Using the latest GWAS datasets, there were 477 (403), 352 (314) and 882 (716) detected by mBAT-combo (mBAT) only (absence of single SNP associations in gene region ± 1Mb and not detected by either fastBAT or MAGMA) for height, BMI and SCZ respectively. We predict that most of these genes, if not all, will be detected in GWAS in the future.

**Figure 6.**
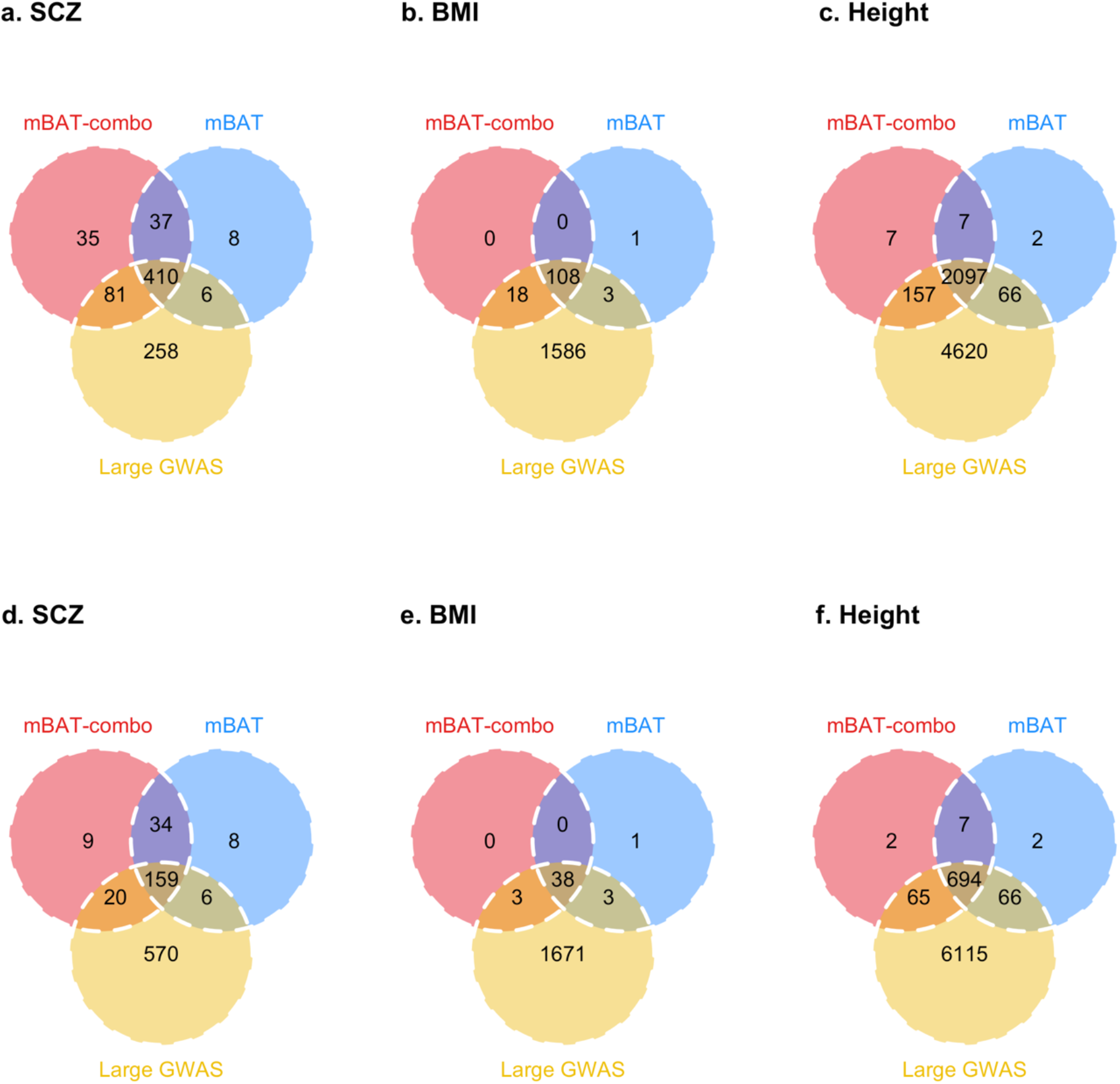
The overlap between the mBAT-combo or mBAT specific genes identified using an earlier release of GWAS data with a smaller samples size and the latest GWAS data with a larger sample size in different traits. In panels a, b, and c, the red (blue) circle shows the number of novel genes detected by mBAT-combo (mBAT) using an earlier GWAS dataset, with no GWAS signal in the gene region ± 1Mb, and the yellow circle shows the numbers of genes with a GWAS signal in the gene region ± 50kb using the latest GWAS dataset and not found in earlier GWAS. In panels d, e and f, the red (blue) circle shows the number of novel gene detected only by mBAT (mBAT-combo) but missed by fastBAT and MAGMA using the earlier GWAS dataset with a smaller sample size.

For example, *LRRC4B*, a susceptibility gene reported by the latest SCZ GWAS study^46^ using FINEMAP^48^, a statistical fine-mapping approach, was detected only by mBAT in this study. *LRRC4B* has been annotated by SynGO^49^, an evidence-based synaptic annotation database, that its protein product plays a role in synaptic organization and differentiation.

## Discussion

In this study, we have introduced two new multivariate gene-based methods, mBAT and mBAT-combo. Briefly, mBAT tests for the nonzero variance explained by all the SNPs in a gene region, using GWAS summary statistics and reference LD. We have shown by theory, simulations, and real trait analyses that mBAT is equivalent to the multiple linear regression approach without access to the individual-level data and improves power of identifying genes harbouring variants with masking effects (when the product of SNP effects and LD correlation is negative). While mBAT was more powerful than the sum-*χ*^2^ methods in the context of masking effects, we observed a small loss of power when genes have multiple causally associated variants have the same direction of effect and the SNPs are in positive LD. Since different genes may have different genetic architectures of association, we introduced the mBAT-combo method that combines the mBAT and fastBAT associations.

The same genes can have different genetic architectures in different traits, which could make them masked in some traits but not in others. This has been confirmed in our analysis of the 35 UKB blood and urine traits. There were 1,483 genes showing multiple association signals in more than 2 traits, among which 579 genes showed both feature of masking and non-masking in different traits according to 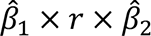 of the top two SNPs reported by COJO. For instance, gene *A1CF* showed a signal of masking effect in creatinine and eGFR marker but a signal of non-masking effect in gamma glutamyl transferase marker. Variants in *ABCA1* were masked in cholesterol adjusted for statins, HDL cholesterol and urate marker but not in apolipoprotein A and LDL direct adjusted for statins. To maximise the overall power, it is necessary to use the mBAT-combo test that accounts for both masking and non-masking effects.

The superiority of the mBAT-combo method over other gene-based test was evidenced by simulations and real trait analyses using individual-level data and the COJO results. As a result, for the 35 UKB biomarker traits we uncovered 98 genes with significant signals of masking effects that cannot be identified by other gene-based methods. To demonstrate that new discoveries are enriched with masking effects, we conducted analyses to provide validation of the discoveries. Pooled across three complex traits, 96.3% (95.5%) of the mBAT (mBAT-combo) specific genes were successfully validated by the conventional GWAS approach in the latest datasets with larger sample sizes, confirming that mBAT and mBAT-combo are powerful methods that produce consistent and reproducible results.

It remains an open question about how common the masking effect is in the human genome. Comprehensive analysis in all traits of UKB is expected to address this question and will be our next project. Still, our analysis provides some insight into this question by quantifying the proportion of genes that are significant in mBAT but not in the sum-*χ*^2^ methods for a wide range of complex traits. For the 35 blood and urine metabolite traits analyzed in this study, we identified a total of 4,273 gene-trait pairs with P_mBAT_ < 5×10^−8^ but P_fastBAT & MAGMA_ > 2.7×10^−6^ in 34 traits, implicating pervasive masking effects in complex traits. Note that 4.1% (4/98) of these genes did not have any SNP with the marginal association P-value reached the genome-wide significance level, which can therefore contribute to explaining the “missing heritability” problem, that is, a fraction of the total variance explained by the independently genome-wide significant SNPs is masked due to the negative genetic covariance between causal variants^19^. Moreover, the presence of masking effect may not occur completely at random. Instead, it could result from the actions of natural selection that have shaped the genetic variation in complex traits. For example, negative LD between two trait-increasing alleles at different loci can be introduced by directional selection (also known as Bulmer effect^21^ or stabilising selection on the trait (**Fig. S17**). A masking effect could also occur if two negatively correlated trait-associated variants are both under selection, even though the trait itself is not correlated with fitness. Given the mounting evidence of natural selection in the human genome^22,23,25,50^, masking effects are likely to be common. We expect that applications of our methods on a broad range of traits will enhance our understanding of masking effects across the genome, their contribution to missing heritability, and the evolutionary processes that give rise to their occurrences.

Another advantage of mBAT is that its test statistic is analytically tractable such that the computation of P-value is exact and fast. The commonly used sum-*χ*^2^ methods such as VEGAS^15^, MAGMA^17^ and fastBAT^16^ use the same statistical test but differ in the way they calculate p-values. VEGAS is computationally slow as P-values are derived from simulations and fastBAT was shown to match VEGAS results with P-values derived from Saddlepoint approximation^35^. MAGMA uses the Imhof’s method^36^ to generate P-values by numerical integration but reverts to simulation when the numerical integration fails. In addition, the Imhof’s method implemented in MAGMA appears to have a limitation on the precision (**Fig. S18**), resulting in a relatively low power comparing to fastBAT and mBAT (**Fig. 3 and 5**). For genes that pass the genome-wide significance threshold, MAGMA underestimates their P-values, which could be an issue in the subsequent analysis that relies on the P-values of these genes, such as pathway or tissue enrichment analysis.

In contrast to the sum-*χ*^2^ methods that do not account for the directions of z-score associations, mBAT, as a multivariate approach, makes use of both direction and size of each z-score, which potentially could result in mBAT being sensitive to the LD differences between GWAS and reference samples as well as various errors in GWAS summary statistics such as genotyping errors or mistakes in labelling the effect alleles. According to our results, removing LD matrix eigenvalues that account for only a small fraction of LD variance (*γ* = 0.9) appeared to be sufficient to address the LD differences due to sampling variation when using external LD, and would allow the method to tolerate a reasonably high proportion of mislabelled effect alleles in the GWAS summary data. Alternatively, the potentially problematic SNPs can be filtered by applying an external tool prior to the analysis, such as DENTIST^47^, which compares the difference between the observed GWAS test-statistic of a variant and its predicted value (using the neighbouring variants in a sliding window and LD data from a reference panel). While DENTIST is versatile in detecting various errors, it can result in a substantial loss in power for some genes (**Fig. S4b, 19-21**). Thus, we set eigenvalue selection without DENTIST as the default setting for mBAT. However, when a small LD reference dataset was used (1KGP), MAGMA tended to give higher P-values for genes with strong signals than those using in-sample LD (**Fig. S20**). In this case, DENTIST could be useful to reduce the inflation.

A feature of all gene-based tests is that gene boundaries overlap so that SNPs can be annotated to more than one gene. There are 33.1% SNPs (458,158/1,382,625) across the whole genome located in more than one gene (gene region defined as gene ± 50kb). A distribution of the number of genes a SNP is allocated to can be found in **Figure S22**, where the highest number of genes a SNP allocated to is 23. It has been found that genes tend to have similar biological functions when collocated^51–53^, and they are likely to be detected together by gene-based tests because of the overlapping of gene regions and the leakage of association signals due to LD between SNPs at the locus. Thus, it requires caution in downstream analyses such as pathway and tissue/cell type enrichment tests, where the enrichment signal could be inflated because of the inclusion of collocated genes. This is a feature of all gene-based tests, and one solution is to perform conditional or joint tests on multiple genes in neighbourhood^54^.

In conclusion, we proposed a multivariate approach that is superior to the prevailing sum-*χ*^2^ methods in uncovering genes with masking effects, and a combined method to maximise the overall power for gene mapping. As a more powerful gene-based method, mBAT or mBAT-combo is expected to improve the downstream pathway analysis or tissue and cell-type enrichment analysis that takes genes identified from GWAS data as input to understand the biological mechanisms of the trait or disease^55^.

## Acknowledgements

This research was supported by the Australian National Health and Medical Research Council (1177268, 1113400, 1173790), and the Westlake Education Foundation. Additional support via US NIMH R01 MH121545. This study makes use of data from dbGaP (accession: phs000788) and the UK Biobank (project ID: 12505). A full list of acknowledgements to these data sets can be found in the Supplementary Note.

## Author contributions

J.Y., J.Z. and N.R.W. conceived and supervised the study. A.L., J.Z., J.Y. and N.R.W. developed the methods and designed the experiment. A.L. conducted all analyses and developed the software tool with the assistance or guidance from A.B., L.J., W.C., Z.Z., P.F.S., P.M.V., N.R.W., J.Y. and J.Z. S.L. implemented GCTA version with assistance of A.L. A.L., J.Z., N.R.W. and J.Y. wrote the manuscript with the participation of all authors. All the authors approved the final version of the manuscript.

## References

1. Locke, A. E. et al. Genetic studies of body mass index yield new insights for obesity biology. Nat. 2015 5187538 518, 197–206 (2015).

2. Visscher, P. M. et al. 10 Years of GWAS Discovery: Biology, Function, and Translation. Am. J. Hum. Genet. 101, 5–22 (2017).

3. Neale, B. M. & Sham, P. C. The future of association studies: Gene-based analysis and replication. Am. J. Hum. Genet. 75, 353–362 (2004).

4. Yang, J. et al. Conditional and joint multiple-SNP analysis of GWAS summary statistics identifies additional variants influencing complex traits. Nat. Genet. 44, 369–375 (2012).

5. Yengo, L. et al. Meta-analysis of genome-wide association studies for height and body mass index in ∼700000 individuals of European ancestry. Hum. Mol. Genet. 27, 3641–3649 (2018).

6. Abell, N. S. et al. Multiple causal variants underlie genetic associations in humans. Science (80-.). 375, 1247–1254 (2022).

7. CA Leeuw, B. N. T. H. D. P. The statistical properties of gene-set analysis. Nat. Rev. Genet. 17, 353–364 (2016).

8. Julienne, H. et al. Multitrait genetic-phenotype associations to connect disease variants and biological mechanisms. 1–25 (2020) doi:10.1101/2020.06.26.172999.

9. Lamparter, D., Marbach, D., Rueedi, R., Kutalik, Z. & Bergmann, S. Fast and Rigorous Computation of Gene and Pathway Scores from SNP-Based Summary Statistics. PLoS Comput. Biol. 12, 1–20 (2016).

10. Gerring, Z. F., Mina-Vargas, A., Gamazon, E. R. & Derks, E. M. E-MAGMA: an eQTL-informed method to identify risk genes using genome-wide association study summary statistics. Bioinformatics (2021) doi:10.1093/BIOINFORMATICS/BTAB115.

11. Watanabe, K., Taskesen, E., Van Bochoven, A. & Posthuma, D. Functional mapping and annotation of genetic associations with FUMA. Nat. Commun. 8, (2017).

12. Skene, N. G. et al. Genetic identification of brain cell types underlying schizophrenia. Nat. Genet. 2018 506 50, 825–833 (2018).

13. Bryois, J. et al. Genetic identification of cell types underlying brain complex traits yields insights into the etiology of Parkinson’s disease. Nat. Genet. 2020 525 52, 482–493 (2020).

14. Martin, A. R. et al. Clinical use of current polygenic risk scores may exacerbate health disparities. Nat. Genet. 2019 514 51, 584–591 (2019).

15. Liu, J. Z. et al. A versatile gene-based test for genome-wide association studies. Am. J. Hum. Genet. 87, 139–145 (2010).

16. Bakshi, A. et al. Fast set-based association analysis using summary data from GWAS identifies novel gene loci for human complex traits. Sci. Rep. 6, 1–9 (2016).

17. de Leeuw, C. A., Mooij, J. M., Heskes, T. & Posthuma, D. MAGMA: Generalized Gene-Set Analysis of GWAS Data. PLoS Comput. Biol. 11, (2015).

18. Yang, J. Conditional and joint multiple-SNP analysis of GWAS summary statistics identifies additional variants influencing complex traits. Nat. Genet. 44, 369–375 (2012).

19. Brown, B. C., Price, A. L., Patsopoulos, N. A. & Zaitlen, N. Local joint testing improves power and identifies hidden heritability in association studies. Genetics 203, 1105–1116 (2016).

20. Sanjak, J. S., Sidorenko, J., Robinson, M. R., Thornton, K. R. & Visscher, P. M. Evidence of directional and stabilizing selection in contemporary humans. Proc. Natl. Acad. Sci. U. S. A. 115, 151–156 (2018).

21. Bulmer, M. G. The Effect of Selection on Genetic Variability. https://doi.org/10.1086/282718 105, 201–211 (2015).

22. Zeng, J. et al. Widespread signatures of natural selection across human complex traits and functional genomic categories. Nat. Commun. 2021 121 12, 1–12 (2021).

23. O’Connor, L. J. et al. Extreme Polygenicity of Complex Traits Is Explained by Negative Selection. Am. J. Hum. Genet. 105, 456–476 (2019).

24. Gazal, S. et al. Functional architecture of low-frequency variants highlights strength of negative selection across coding and non-coding annotations. Nat. Genet. 2018 5011 50, 1600–1607 (2018).

25. Zeng, J. et al. Signatures of negative selection in the genetic architecture of human complex traits. Nat. Genet. 2018 505 50, 746–753 (2018).

26. Shi, H., Kichaev, G. & Pasaniuc, B. Contrasting the Genetic Architecture of 30 Complex Traits from Summary Association Data. Am. J. Hum. Genet. 99, 139–153 (2016).

27. Ehret, G. B. et al. A multi-SNP locus-association method reveals a substantial fraction of the missing heritability. Am. J. Hum. Genet. 91, 863–871 (2012).

28. Manolio, T. A. et al. Finding the missing heritability of complex diseases. Nature 461, 747–53 (2009).

29. Eichler, E. E. et al. Missing heritability and strategies for finding the underlying causes of complex disease. Nat. Rev. Genet. 2010 116 11, 446–450 (2010).

30. Wainschtein, P. et al. Assessing the contribution of rare variants to complex trait heritability from whole-genome sequence data. Nat. Genet. 2022 1–11 (2022) doi:10.1038/s41588-021-00997-7.

31. Ferrari, A. A note on sum and difference of correlated chi-squared variables. (2019).

32. Sinnott-Armstrong, N. et al. Genetics of 35 blood and urine biomarkers in the UK Biobank. Nat. Genet. 2021 532 53, 185–194 (2021).

33. Yang, J., Lee, S. H., Goddard, M. E. & Visscher, P. M. GCTA: A tool for genome-wide complex trait analysis. Am. J. Hum. Genet. 88, 76–82 (2011).

34. Enderlein, G. Scheffé, H.: The Analysis of Variance. Wiley, New York 1959, 477 Seiten, $ 14,00. Biom. Z. 3, 143–144 (1961).

35. Kuonen, D. Saddlepoint approximations for distributions of quadratic forms in normal variables. Biometrika 86, (1999).

36. P.J., I. Computing the distribution of quadratic forms in normal variables. Biometrika 48, 419 (1961).

37. Liu, Y. et al. ACAT: A Fast and Powerful p Value Combination Method for Rare-Variant Analysis in Sequencing Studies. Am. J. Hum. Genet. 104, 410–421 (2019).

38. Liu, Y. & Xie, J. Cauchy Combination Test: A Powerful Test With Analytic p-Value Calculation Under Arbitrary Dependency Structures. J. Am. Stat. Assoc. 115, 393–402 (2020).

39. Ha, H. T. & Provost, S. B. An accurate approximation to the distribution of a linear combination of non-centralchi-squarerandomvariables. Revstat Stat. J. 11, 231–254 (2013).

40. Kvale, M. N. et al. Genotyping informatics and quality control for 100,000 subjects in the genetic epidemiology research on adult health and aging (GERA) cohort. Genetics 200, 1051–1060 (2015).

41. Altshuler, D. M. et al. An integrated map of genetic variation from 1,092 human genomes. Nat. 2012 4917422 491, 56–65 (2012).

42. Jiang, L. et al. A resource-efficient tool for mixed model association analysis of large-scale data. Nat. Genet. 2019 5112 51, 1749–1755 (2019).

43. Wood, A. R. et al. Defining the role of common variation in the genomic and biological architecture of adult human height. Nat. Genet. 2014 4611 46, 1173–1186 (2014).

44. Locke, A. E. et al. Genetic studies of body mass index yield new insights for obesity biology. Nature 518, 197–206 (2015).

45. Ripke, S. et al. Biological insights from 108 schizophrenia-associated genetic loci. Nat. 2014 5117510 511, 421–427 (2014).

46. Trubetskoy, V. et al. Mapping genomic loci implicates genes and synaptic biology in schizophrenia. Nat. 2022 6047906 604, 502–508 (2022).

47. Chen, W. et al. Improved analyses of GWAS summary statistics by reducing data heterogeneity and errors. Nat. Commun. 2021 121 12, 1–10 (2021).

48. Benner, C. et al. FINEMAP: Efficient variable selection using summary data from genome-wide association studies. Bioinformatics 32, (2016).

49. Koopmans, F. et al. SynGO: An Evidence-Based, Expert-Curated Knowledge Base for the Synapse. Neuron 103, 217-234.e4 (2019).

50. Gazal, S. et al. Linkage disequilibrium-dependent architecture of human complex traits shows action of negative selection. Nat. Genet. 49, 1421–1427 (2017).

51. Oliver, B. & Misteli, T. A non-random walk through the genome. Genome Biol. 6, 1–6 (2005).

52. De, S. & Babu, M. M. Genomic neighbourhood and the regulation of gene expression. Curr. Opin. Cell Biol. 22, 326–333 (2010).

53. Emmert-Streib, F. et al. Functional and genetic analysis of the colon cancer network. BMC Bioinformatics 15, 1–15 (2014).

54. Mancuso, N. et al. Probabilistic fine-mapping of transcriptome-wide association studies. Nat. Genet. 2019 514 51, 675–682 (2019).

55. Quick, C., Wen, X., Abecasis, G., Boehnke, M. & Kang, H. M. Integrating comprehensive functional annotations to boost power and accuracy in gene-based association analysis. PLOS Genet. 16, e1009060 (2020).

