## Supplementary Note for "mBAT-combo: a more powerful test to detect gene-trait associations from GWAS data"

#### Distribution of mBAT test statistic

Let $\mathbf{z}$ be a vector of GWAS z-scores of SNPs in the target gene region, with $z_j = \frac{b_j}{SE_j}$ where $b_j$ is the marginal effect estimate of SNP $j$ and $SE_j$ is the standard error. Assuming both genotypes ($\mathbf{X}_j$) and phenotypes ($\boldsymbol{y}$) are standardised with mean zero and unit variance, $b_j = (\mathbf{X}_j'\mathbf{X}_j)^{-1}\mathbf{X}_j'\mathbf{y} = \frac{1}{\mathbf{n}}\mathbf{X}_j'\mathbf{y}$, with $\mathbf{n}$ being the sample size, and $SE_j = \sqrt{(\mathbf{X}_j'\mathbf{X}_j)^{-1}\hat{\sigma}_e^2} = \sqrt{\frac{1}{\mathrm{n}}\left(\sigma_y^2 - \sigma_{X_j}^2 b_j^2\right)} \approx \frac{1}{\sqrt{\mathrm{n}}}$, provided that negligible variance is explained by a single SNP ($b_j^2 \approx 0$). Therefore,

$$\mathrm{E}[\mathbf{z}] = \frac{1}{\sqrt{\mathrm{n}}}\mathbf{X}'\mathbf{y} = \boldsymbol{\mu}$$

$$\mathrm{Var}[\mathbf{z}] = \frac{1}{\mathrm{n}}\mathbf{X}'\mathrm{Var}[\mathbf{y}]\mathbf{X} = \frac{1}{\mathrm{n}}\mathbf{X}'\mathbf{X} = \mathbf{R}$$

where $\mathbf{R}$ is the LD correlation matrix among SNPs that can be obtained from a reference sample. Assuming normality, the sampling distribution of $\mathbf{z}$ is

$$\mathbf{z} \sim N(\boldsymbol{\mu}, \mathbf{R})$$

The test statistic of mBAT is

$$T_{\mathrm{mBAT}} = \mathbf{z}'\mathbf{R}^{-1}\mathbf{z}$$

Let $\boldsymbol{\Lambda}$ and $\mathbf{U}$ be the diagonal matrix of eigenvalues and the orthogonal matrix of eigenvectors of $\mathbf{R}$, respectively. We have

$$\mathbf{R}^{-1} = \mathbf{U}\boldsymbol{\Lambda}^{-1}\mathbf{U}'$$

Let $\mathbf{w} = \boldsymbol{\Lambda}^{-\frac{1}{2}}\mathbf{U}'\mathbf{z}$. Then,

$$T_{\mathrm{mBAT}} = \mathbf{z}'\mathbf{U}\boldsymbol{\Lambda}^{-1}\mathbf{U}'\mathbf{z} = \mathbf{w}'\mathbf{w}$$

Now consider the distribution of $\mathbf{w}$. Since $\mathrm{E}[\mathbf{w}] = \boldsymbol{\Lambda}^{-\frac{1}{2}}\mathbf{U}'\boldsymbol{\mu}$ and $\mathrm{Var}[\mathbf{w}] = \boldsymbol{\Lambda}^{-\frac{1}{2}}\mathbf{U}'\mathbf{R}\mathbf{U}\boldsymbol{\Lambda}^{-\frac{1}{2}} = \boldsymbol{\Lambda}^{-\frac{1}{2}}\mathbf{U}'\mathbf{U}\boldsymbol{\Lambda}\mathbf{U}'\mathbf{U}\boldsymbol{\Lambda}^{-\frac{1}{2}} = \mathbf{I}$, it is a normal distribution

$$\mathbf{w} \sim N\left(\boldsymbol{\Lambda}^{-\frac{1}{2}}\mathbf{U}'\boldsymbol{\mu}, \mathbf{I}\right)$$

Under the null hypothesis of no SNP-trait association, $\boldsymbol{\mu} = \mathbf{0}$. In this case, $\mathbf{w}$ is a $\mathrm{m} \times 1$ vector of standard normal variables ($m$ is the number of SNPs). Thus, the test statistic follows a central chi-squared distribution with $m$ degrees of freedom.

$$\text{Under } \mathrm{H}_0\text{:} \quad T_{\mathrm{mBAT}} \sim \chi_m^2$$

Under the alternative hypothesis that there exists SNP-trait associations, $\boldsymbol{\mu} \neq \mathbf{0}$. Let $\boldsymbol{\eta} = \boldsymbol{\Lambda}^{-\frac{1}{2}}\mathbf{U}'\boldsymbol{\mu}$. Then, $\mathbf{w} \sim N(\boldsymbol{\eta}, \mathbf{I})$. The sum of squares of $\mathbf{w}$ follows a non-central chi-squared distribution with $m$ degrees of freedom and a non-central parameter $NCP = \boldsymbol{\eta}'\boldsymbol{\eta} = \boldsymbol{\mu}'\mathbf{U}\boldsymbol{\Lambda}^{-\frac{1}{2}}\boldsymbol{\Lambda}^{-\frac{1}{2}}\mathbf{U}'\boldsymbol{\mu} = \boldsymbol{\mu}'\mathbf{U}\boldsymbol{\Lambda}^{-1}\mathbf{U}'\boldsymbol{\mu} = \boldsymbol{\mu}'\mathbf{R}^{-1}\boldsymbol{\mu}$.

$$\text{Under H}_1\text{:}\quad T_{\text{mBAT}} \sim \chi^2_m(\boldsymbol{\mu}'\mathbf{R}^{-1}\boldsymbol{\mu})$$

When the pseudo-inverse of **R** (denoted as $\mathbf{R}^-$) is used by keeping the top $k$ eigenvalues or eigenvectors that cumulatively explain $\gamma$ proportion of variance in LD, i.e., $\gamma = \sum_{j=1}^{k} \Lambda_j \,/ \sum_{j=1}^{m} \Lambda_j$, $\mathbf{w}$ becomes a $k \times 1$ vector of standard normal variables under the null, and a $k \times 1$ vector of normal variables with nonzero means under the alternative ($\mathrm{E}[\mathbf{w}] = \boldsymbol{\Lambda}_{(k)}^{-\frac{1}{2}}\mathbf{U}'_{(k)}\boldsymbol{\mu}$ with $\boldsymbol{\Lambda}_{(k)}^{-\frac{1}{2}}$ being a $k \times k$ diagonal matrix and $\mathbf{U}'_{(k)}$ being a $k \times m$ matrix). As a result,

$$\text{Under H}_0\text{:}\quad T_{\text{mBAT}} \sim \chi^2_k$$

$$\text{Under H}_1\text{:}\quad T_{\text{mBAT}} \sim \chi^2_k(\boldsymbol{\mu}'\mathbf{R}^{-}\boldsymbol{\mu})$$


### Acknowledgements

#### UKB

This study has been conducted using UK Biobank resource under Application Number 12505. UK Biobank was established by the Wellcome Trust medical charity, Medical Research Council, Department of Health, Scottish Government and the Northwest Regional Development Agency. It has also had funding from the Welsh Assembly Government, British Heart Foundation and Diabetes UK.

#### GERA

The Genetic Epidemiology Research on Adult Health and Aging study was supported by grant RC2 AG036607 from the National Institutes of Health, grants from the Robert Wood Johnson Foundation, the Ellison Medical Foundation, the Wayne and Gladys Valley Foundation and Kaiser Permanente. The authors thank the Kaiser Permanente Medical Care Plan, Northern California Region (KPNC) members who have generously agreed to participate in the Kaiser Permanente Research Program on Genes, Environment and Health (RPGEH).

**Supplementary Figures**

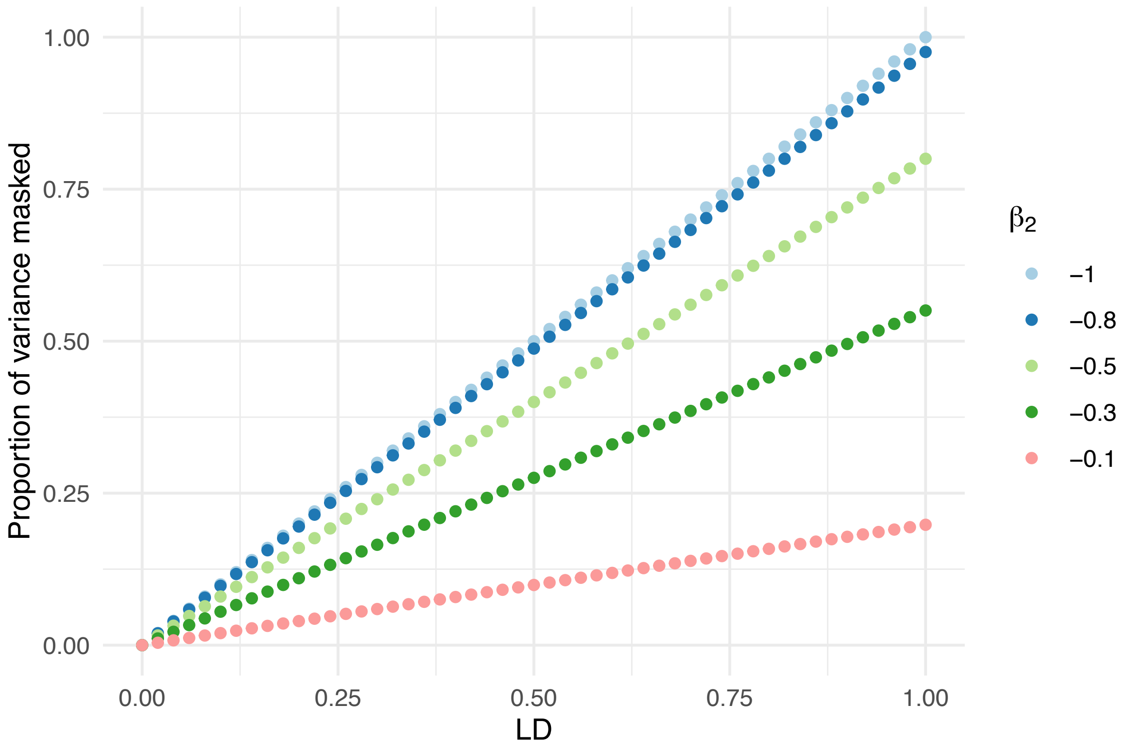


**Supplementary Figure 1** The proportion of variance being masked as a function of LD correlation (*r*) between two variants and the effect size of variant 2 ($\beta_2$), with the effect size of variant 1 $\beta_1 = 1$. The proportion of variance being masked $= \frac{-2\times\beta_1\times\beta_2\times r}{\beta_1^2+\beta_2^2}$.

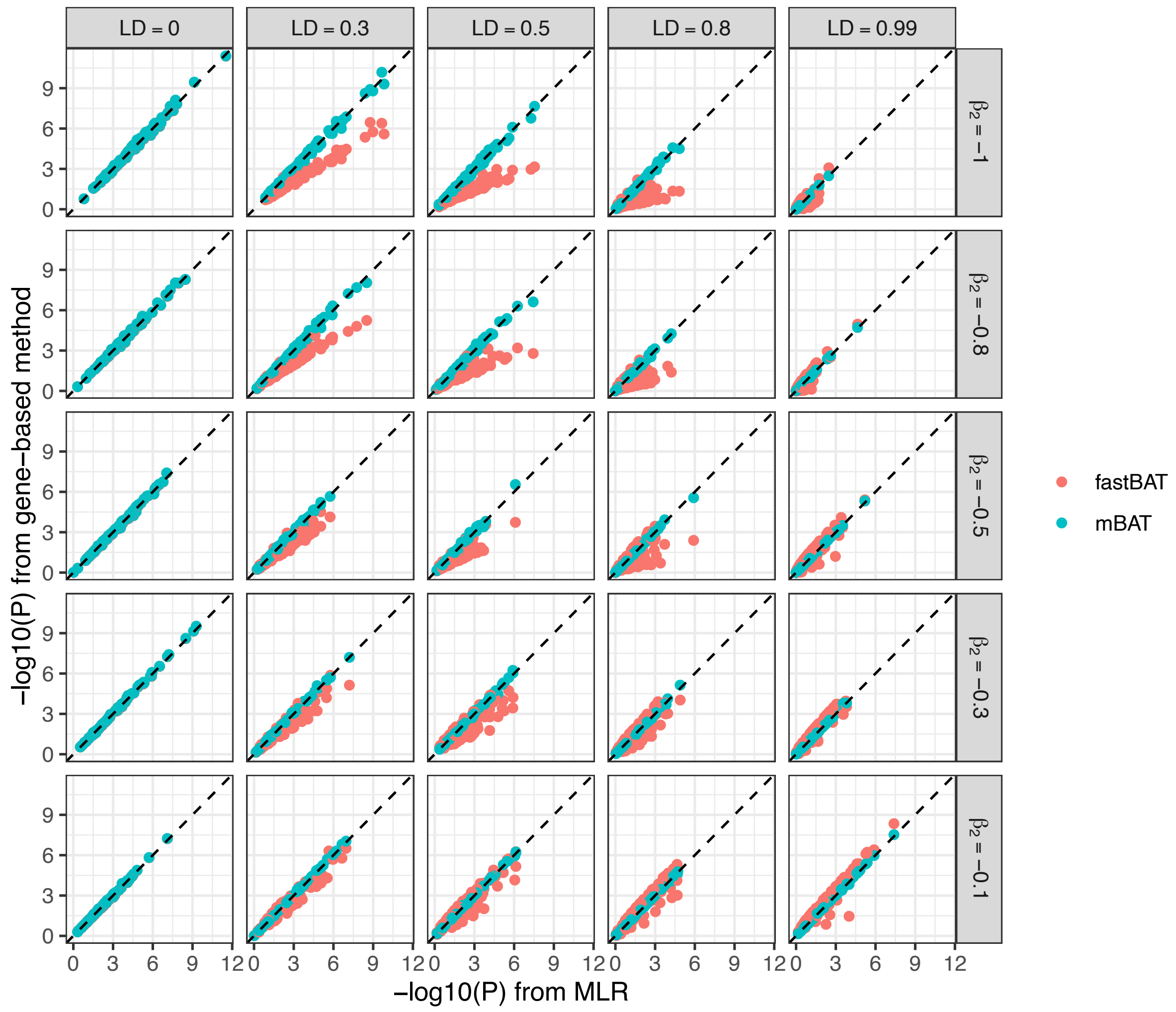


**Supplementary Figure 2** Comparison of -log$_{10}$(*P*-value) from fastBAT or mBAT with that from multiple linear regression (MLR) as the gold standard in the scenario of masking effect. The effect size of variant 1 $\beta_1 = 1$. When LD *r* =0 (left-most plots), the variants were independent, and there was no masking effect. In this case, the results of different methods were almost identical. When LD was very high *r* = 0.99 (right-most plots), the variants were so tightly correlated that the effective number of variants $\approx 1$ and the results from different methods were again similar. For LD correlations between the two extremes, the result of mBAT was in high concordance with that of the gold standard MLR, whereas fastBAT gave less consistent result as the masking effect increased (shown by rows).

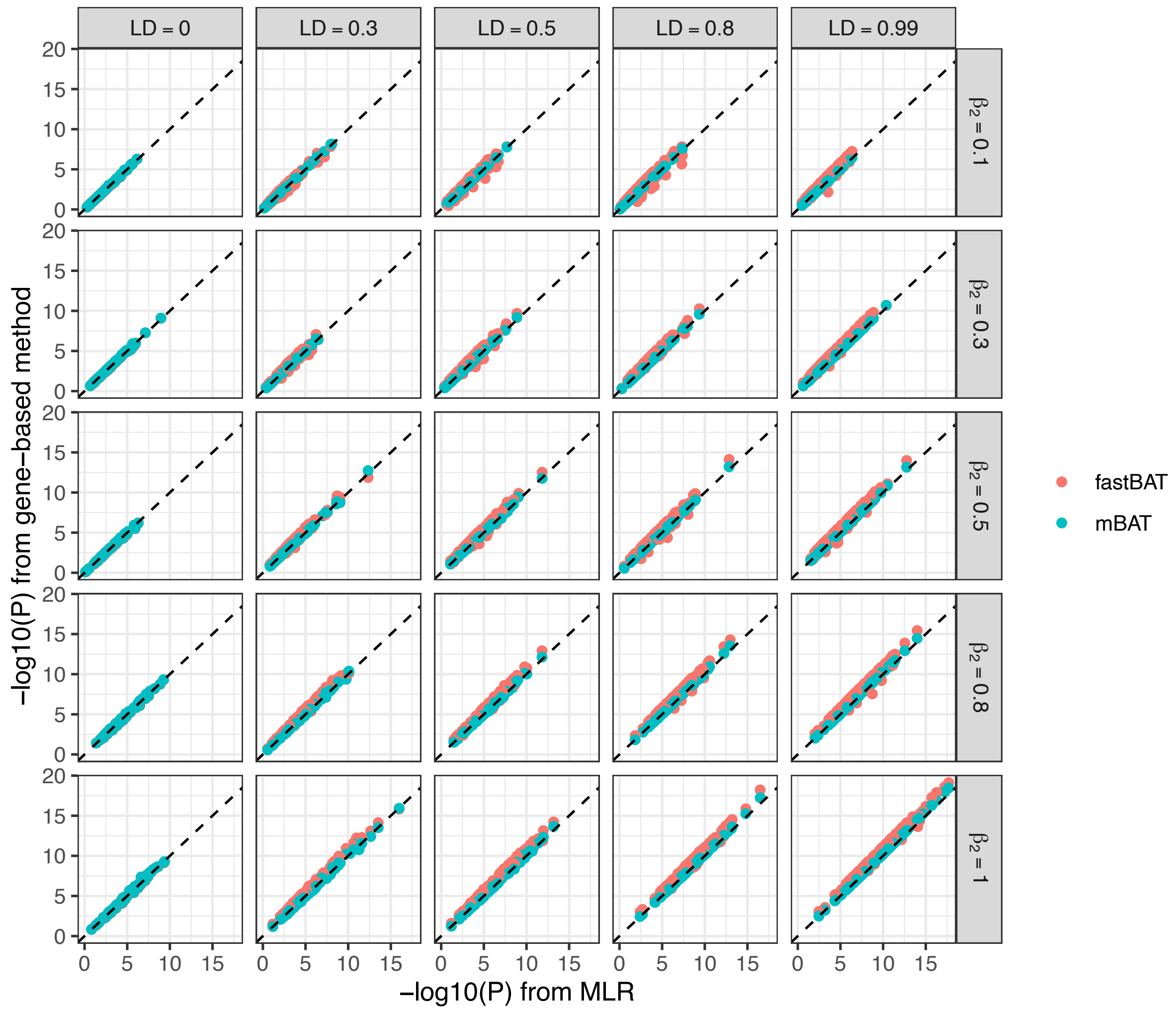


**Supplementary Figure 3** Comparison of -$\log_{10}$(*P*-value) from fastBAT or mBAT with that from multiple linear regression (MLR) as the gold standard in the scenario of non-masking effect. The effect size of variant 1 $\beta_1 = 1$. Regardless of the LD correlation and effect size of variant 2, mBAT gave identical result to MLR.

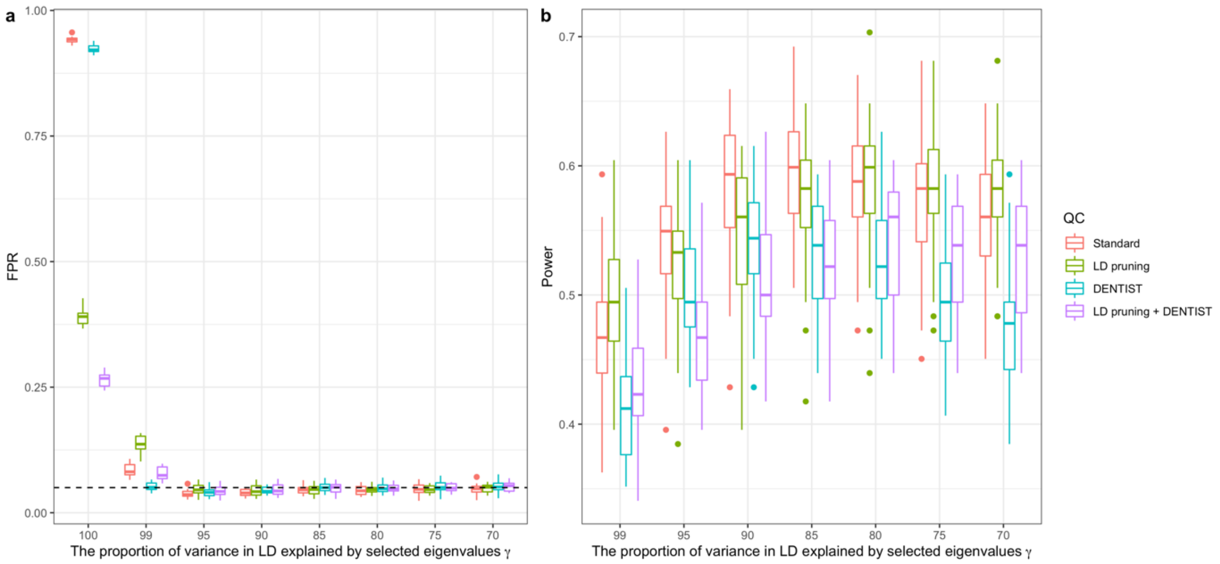


**Supplementary Figure 4** Comparison of the false positive rate and power at different $\gamma$ values for mBAT, with or without additional QC steps. $\gamma$ is the proportion of variance in LD explained by the selected eigenvectors in mBAT. Colours represent different QC steps (Standard: no DENTIST applied nor LD pruning) prior to the mBAT test. a) False positive rate (FPR), quantified from 1,976 genes on chromosome 1 under the null model. Each boxplot represents the distribution of FPR across 25 simulation replicates. b) Power for detecting genes with causal variants, quantified from 90 non-overlapping genes-causal variants pairs (nQTLs= 2~6) on chromosome 1 under the causal model. Each boxplot represents the distribution of power 25 simulation replicates.

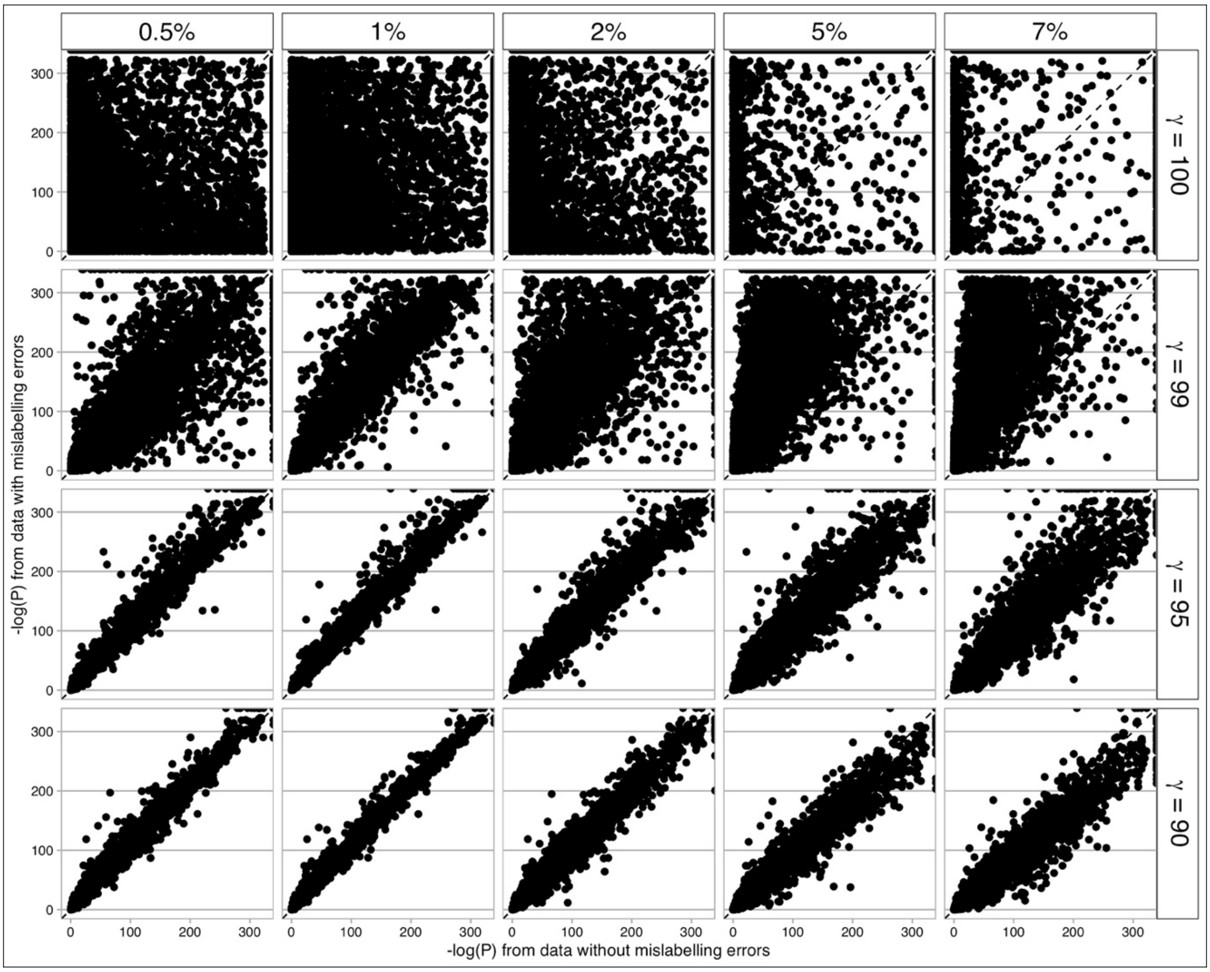


**Supplementary Figure 5** The robustness of mBAT to the allelic mislabelling errors in GWAS data at different $\gamma$ values. The x-axis is the -log10 P-value using GWAS data without errors and in-sample LD (the ideal scenario). The y-axis is the -log10 P-value using GWAS data with a proportion of SNPs whose effect allele was mislabelled. Columns are different proportion of SNPs being mislabelled. Rows are different $\gamma$ values used in mBAT, which is the proportion of variance in LD explained by the selected eigenvectors. The mBAT test was performed without applying additional QC such as DENTIST on the GWAS summary statistics.

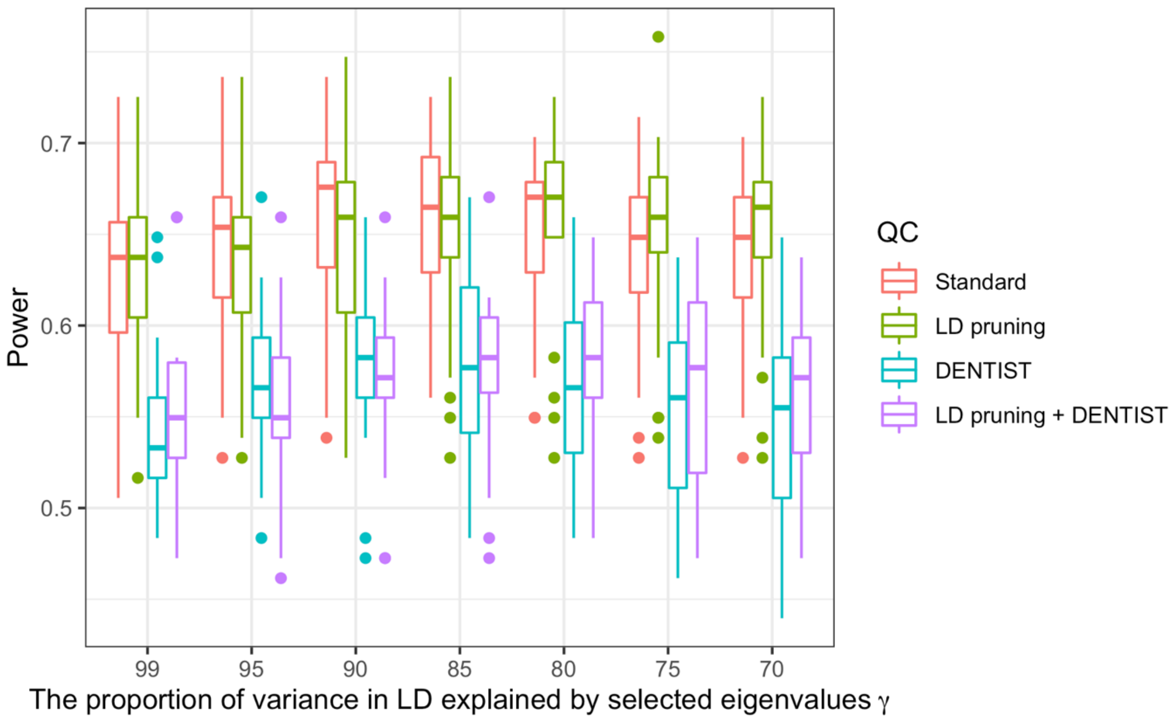


**Supplementary Figure 6** Comparison of the power of mBAT-combo with mBAT at different $\gamma$ values, with or without additional QC steps. $\gamma$ is the proportion of variance in LD explained by the selected eigenvectors in mBAT. Colours represent different QC steps (Standard: no DENTIST applied nor LD pruning) prior to the mBAT test. Power for detecting genes with causal variants, quantified from 90 non-overlapping genes-causal variants pairs (nQTLs= 2~6) on chromosome 1 under the causal model. Each boxplot represents the distribution of power across 25 causal simulation replicates.

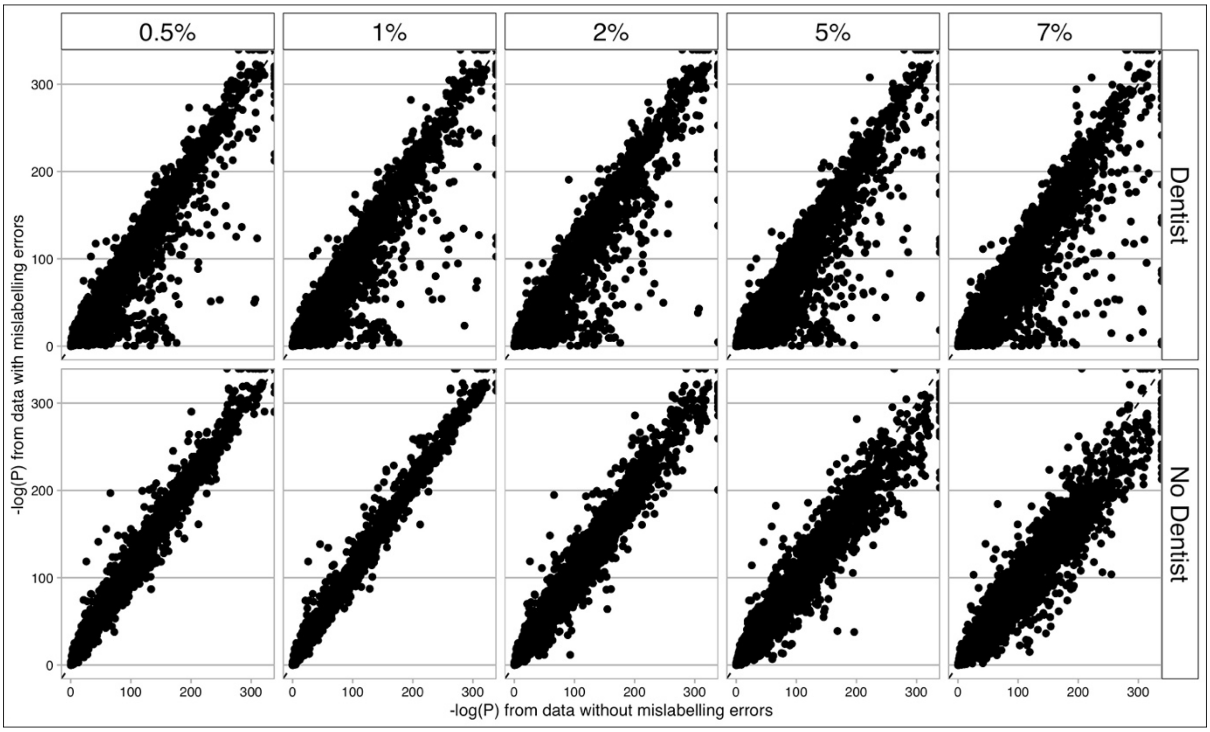


**Supplementary Figure 7** The effect of DENTIST to the allelic mislabelling errors in GWAS data at $\gamma = 90$ set in mBAT. The x-axis is the -log10 P-value using GWAS data without errors and in-sample LD (the ideal scenario). The y-axis is the -log10 P-value using GWAS data with a proportion of SNPs whose effect allele was mislabelled. Columns are different proportion of SNPs being mislabelled. Rows are application of DENTIST QC and no application of DENTIST in mBAT.

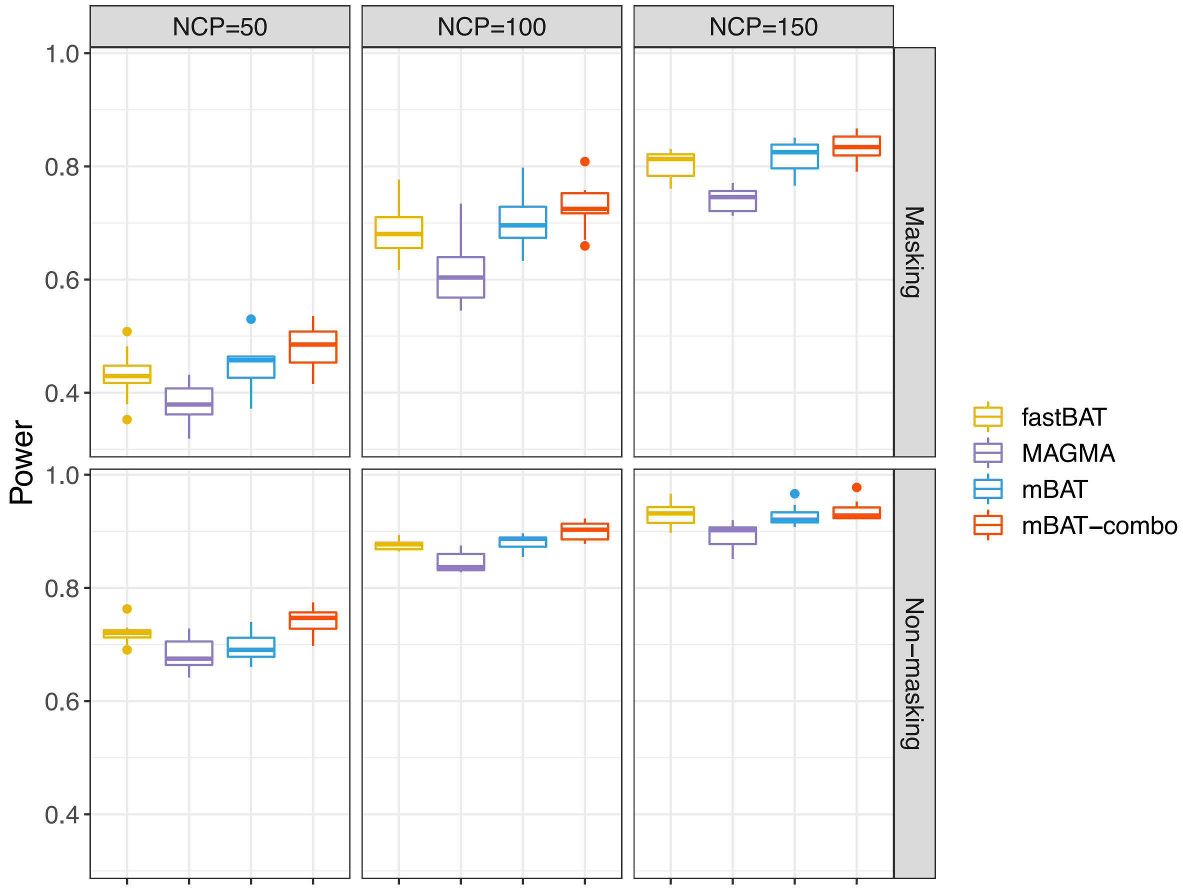


**Supplementary Figure 8**. Power comparison of methods with different NCP levels per gene in scenarios of masking and non-masking effects in the simulation. Each boxplot represents the distribution of estimates across 10 data points

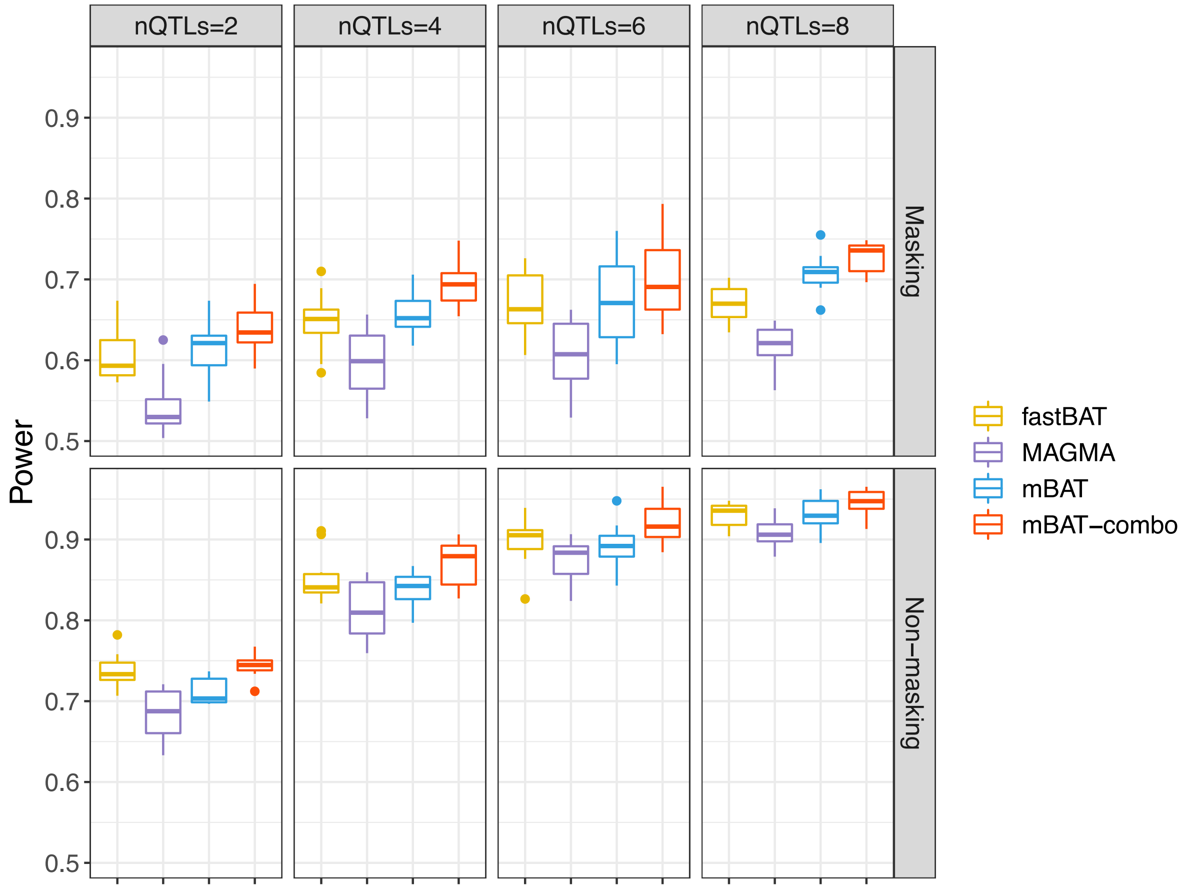


**Supplementary Figure 9** Power comparison of methods with different numbers of causal variants per gene in scenarios of masking and non-masking effects in the simulation. Each boxplot represents the distribution of estimates across 10 data points.

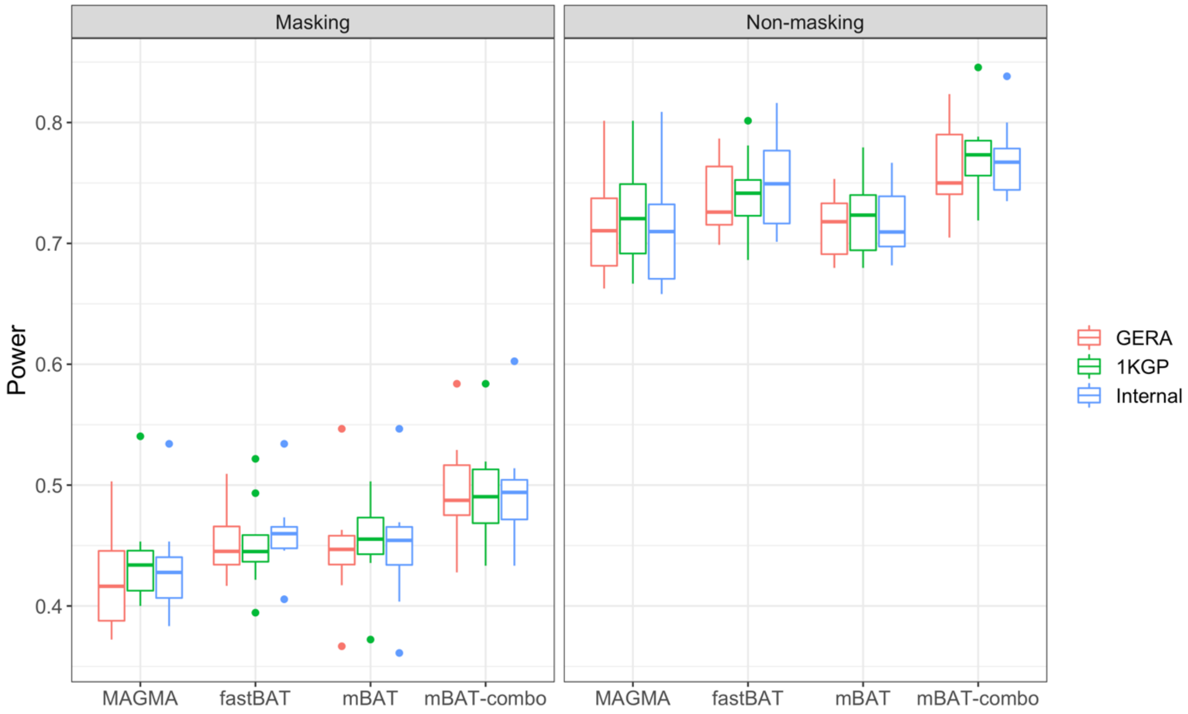


**Supplementary Figure 10** Power comparison of methods with internal (in-sample) or external LD reference in scenarios of masking and non-masking effects in the simulation. Each boxplot represents the distribution of estimates across 10 data points.

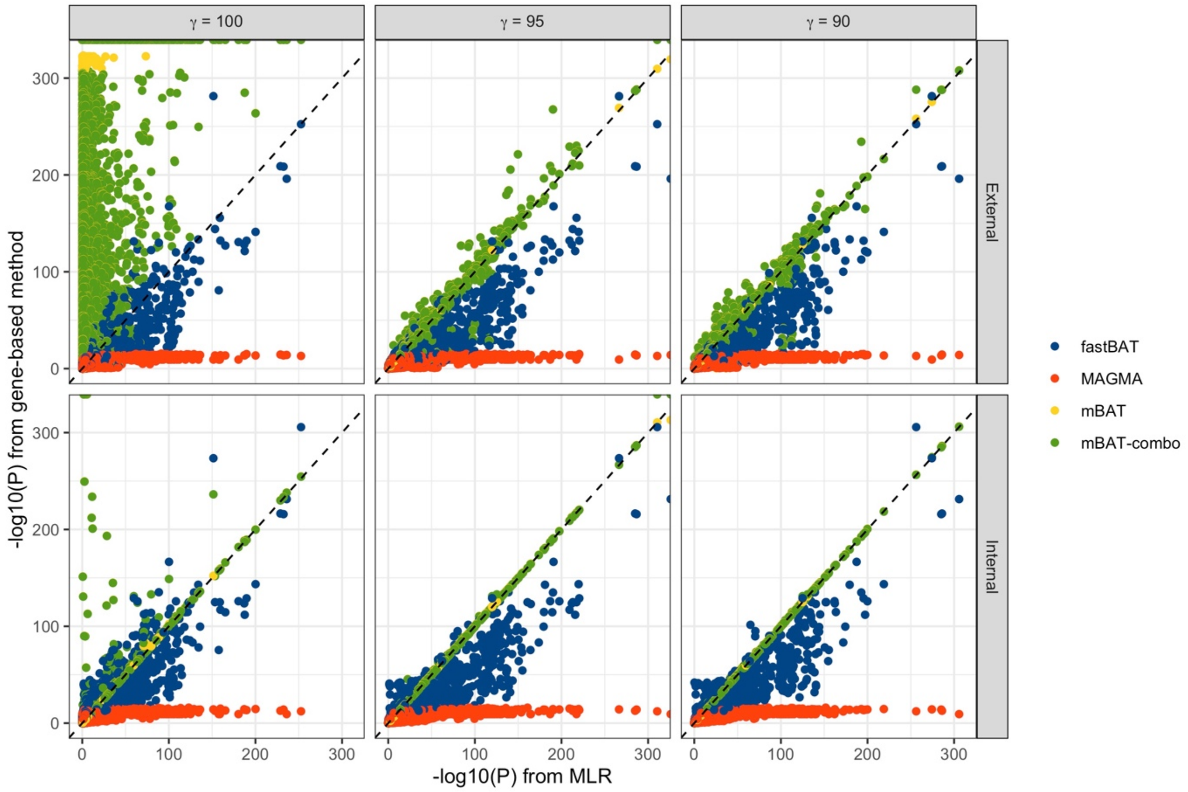


**Supplementary Figure 11** Benchmark of different gene-based methods with multiple linear regression (MLR) in the UKB height, using external LD and in-sample and different $\gamma$ values in mBAT and mBAT-combo. The y-axis is the -log10-P-values of different gene-based methods. The x-axis is the -log10-P-values of MLR with the same number of eigenvectors as that used in mBAT and mBAT-combo.

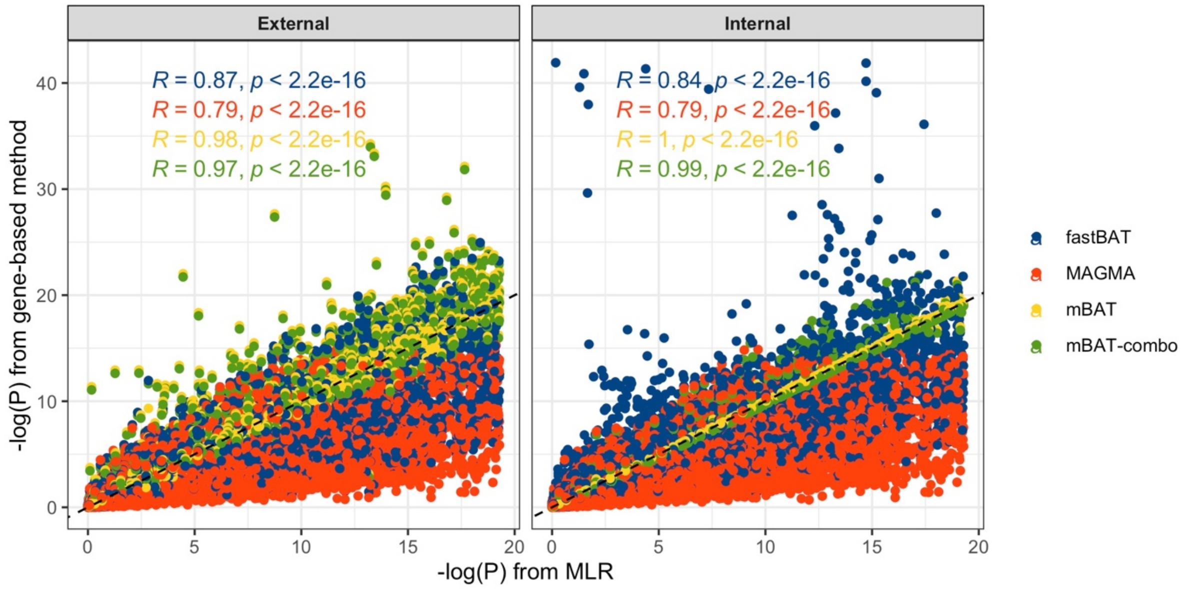


**Supplementary Figure 12** A zoom-in plot of Figure 4 for a benchmark of different gene-based methods with multiple linear regression in the UKB height (the P-value of MLR > $5\times10^{-20}$. The y-axis is the -log10 P-values of different gene-based methods denoted by colours. The x-axis is the -log10 P-values of MLR with the same number of eigenvectors as that used in mBAT and mBAT-combo.

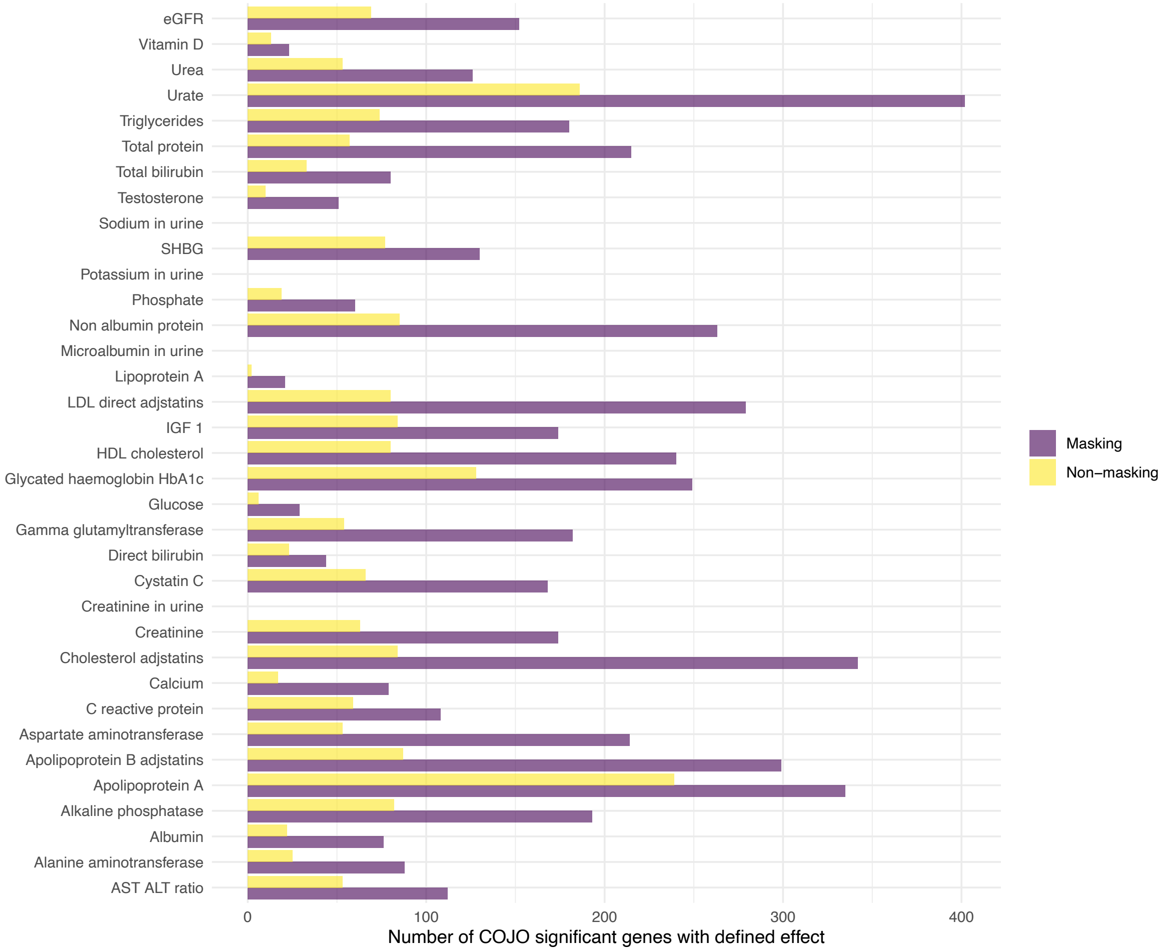


**Supplementary Figure 13** The number of COJO – joint significant genes with masking effect (purple) and non-masking effect (yellow) across 35 blood and urine metabolite biomarkers.

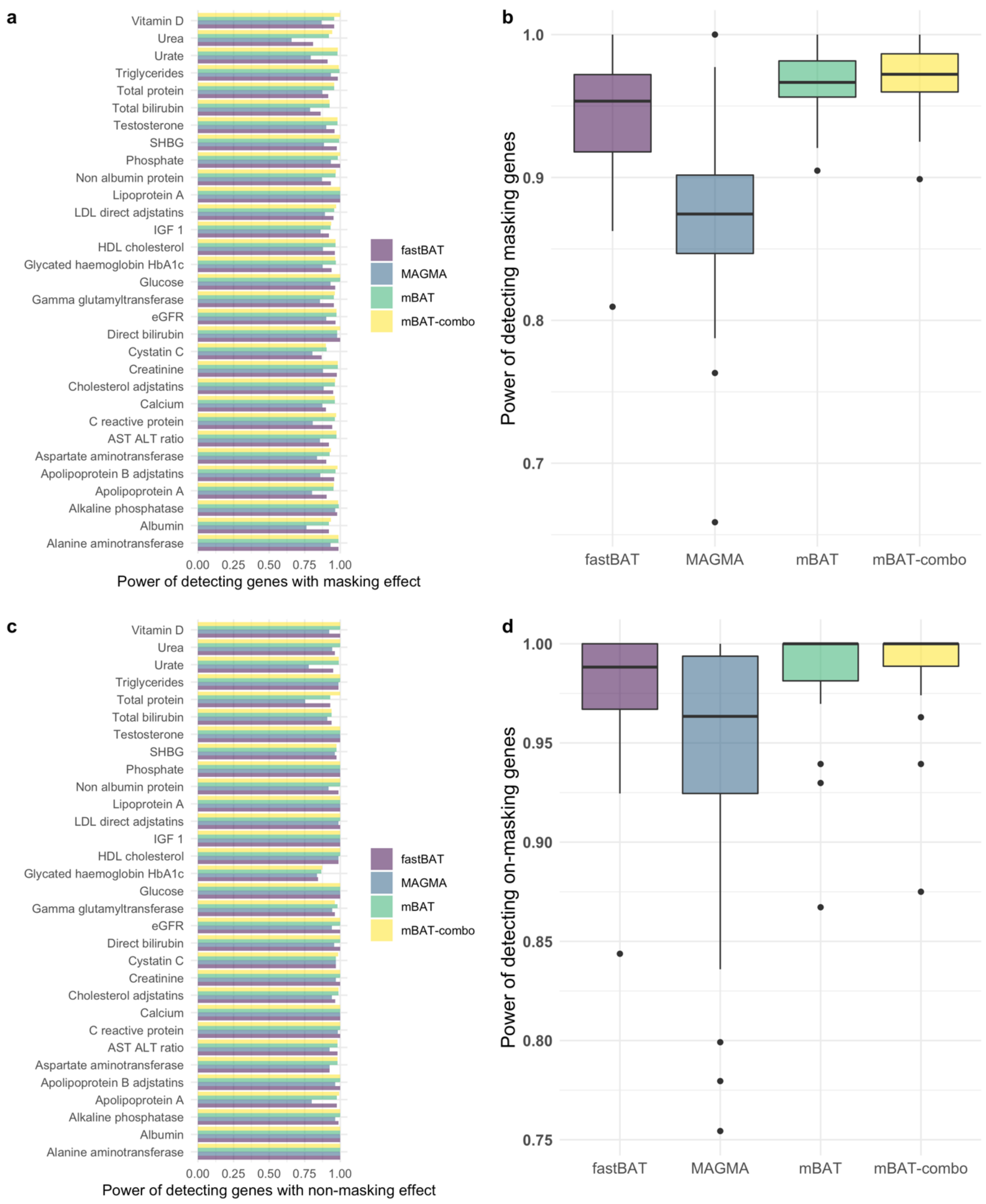


**Supplementary Figure 14** Comparison of power (Panel a&c) and the average power (Panel b&d) across all traits in panel a of detecting genes with masking(a&b) and non-masking (c&d) effects of each of the methods (mBAT-combo, fastBAT, MAGMA, mBAT in different colours) across 35 blood metabolites. Each boxplot represents the distribution of power estimates of each of the methods across 35 blood metabolites. Y-axis is the power of detecting genes with masking/non-masking effects identified by COJO.

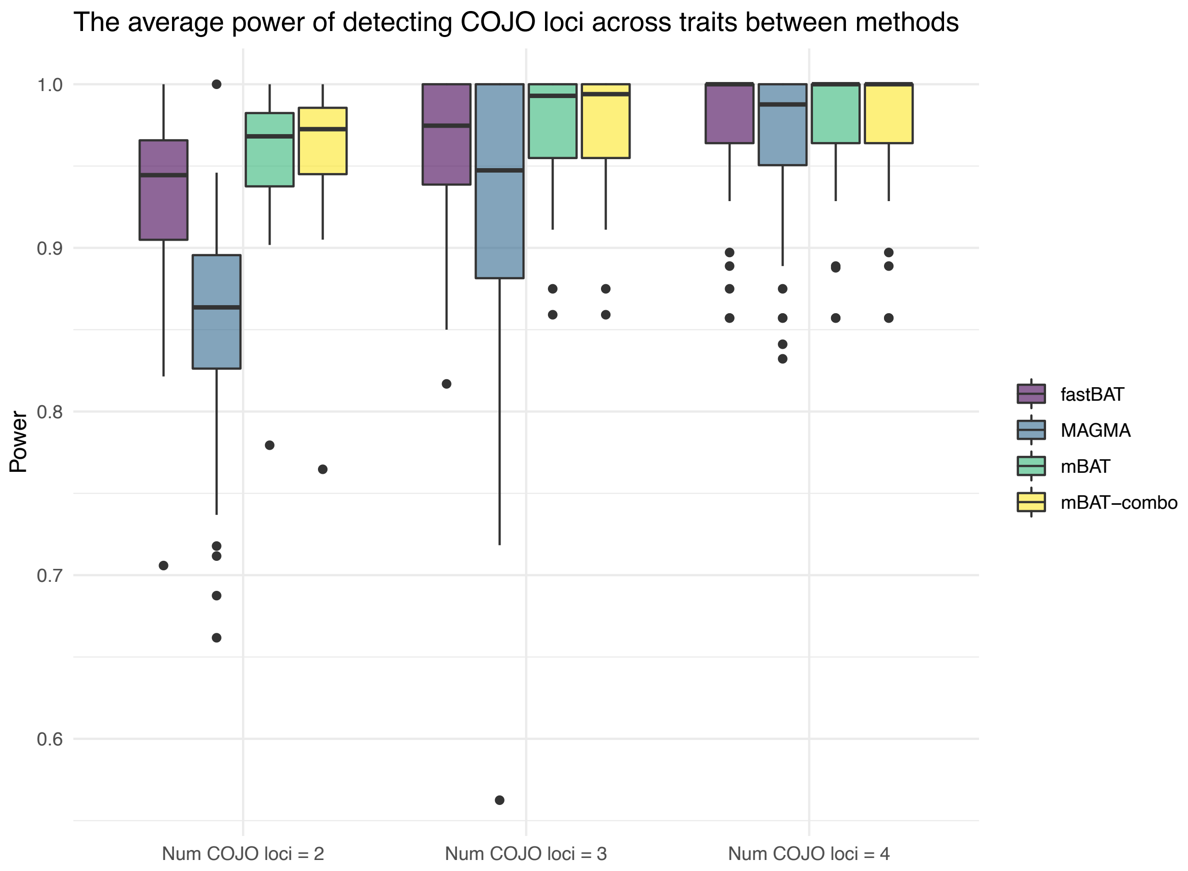


**Supplementary Figure 15** Comparison of power of detecting genes with different numbers of COJO signals between gene-based methods (fastBAT, MAGMA, mBAT and mBAT-combo in different colours) for 35 blood metabolite traits in the UKB.

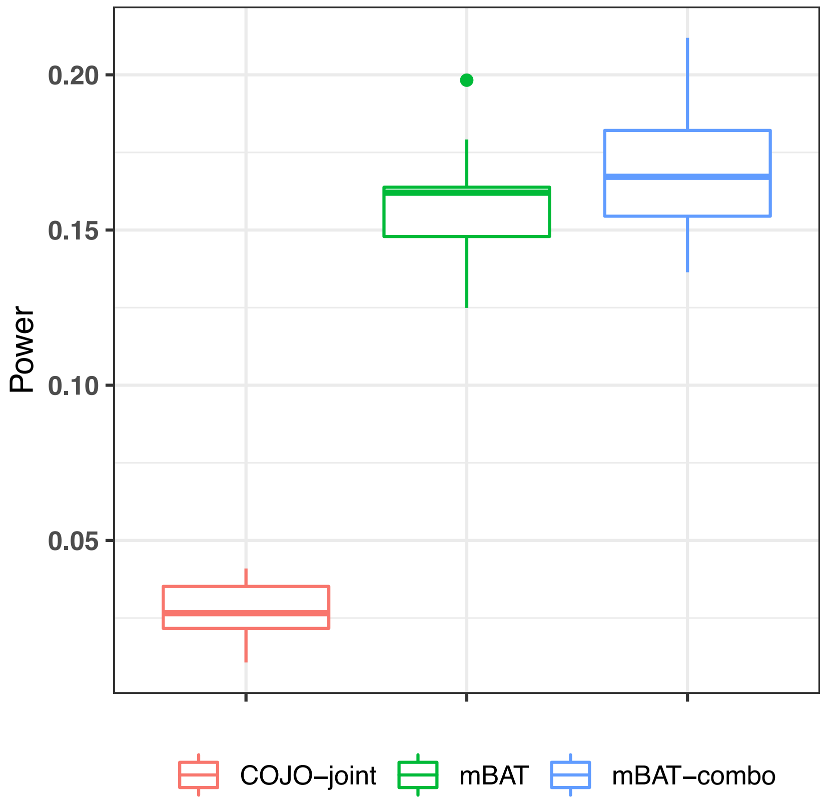


**Supplementary Figure 16.** Power comparison of COJO, mBAT and mBAT-combo for detecting genes with masking effects in simulation. Simulations were based on randomly selecting 2 causal variants per gene among 91 non-overlapping genes on Chromosome 1 with their effect sizes $\beta_j$ sampled from a normal distribution and the directions of the effect sizes assigned in a way that $\beta_1 \times r \times \beta_2$< 0, i.e., a scenario of masking effect. The trait heritability $h^2 = 0.1$. The simulation was carried out for 10 replicates.

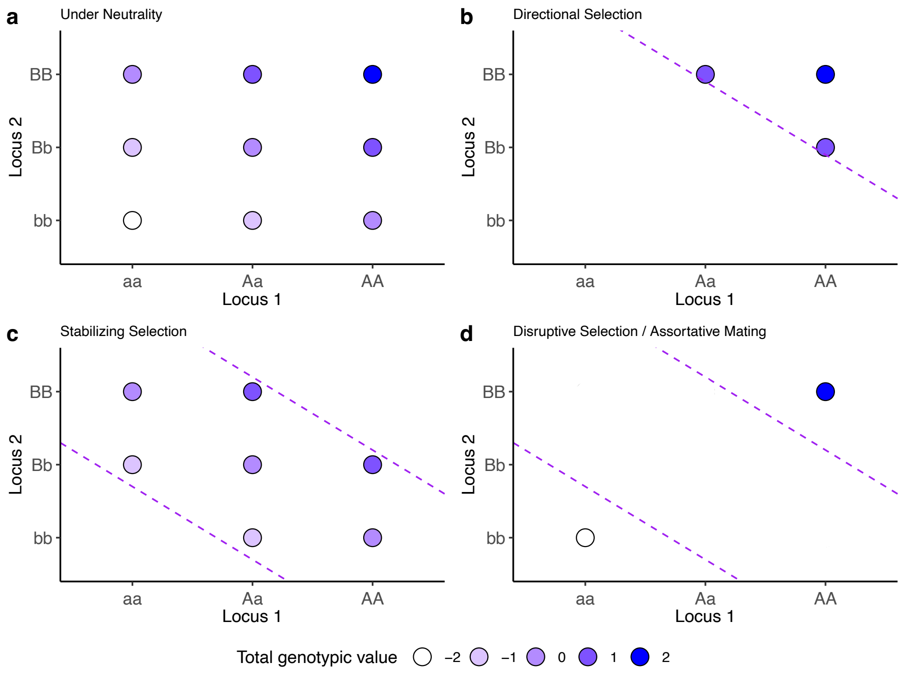


**Supplementary Figure 17.** Schematics showing that different types of selection introduce different directions of LD between two loci. The x- and y-axis are genotypes of two loci: AA, Aa, and aa for locus 1 with genotypic values of 1, 0 and, -1, respectively, and BB, Bb, and bb for locus 2 with the same genotypic values of 1, 0, and -1, respectively. Dots are the individuals who carry the specific combinations of the genotypes with their total genotypic values shown by colour. a) Two independent loci in a population under neutrality; b) directional selection that only keeps individuals with high total genotypic values in the population introduces a negative LD between loci; c) stabilizing selection that keeps individuals with average total genotypic values in the population also introduces a negative LD between loci; and d) disruptive selection that keeps individuals with either high or low total genotypic values introduces a positive LD between loci; this can also be achieved by assortative mating.

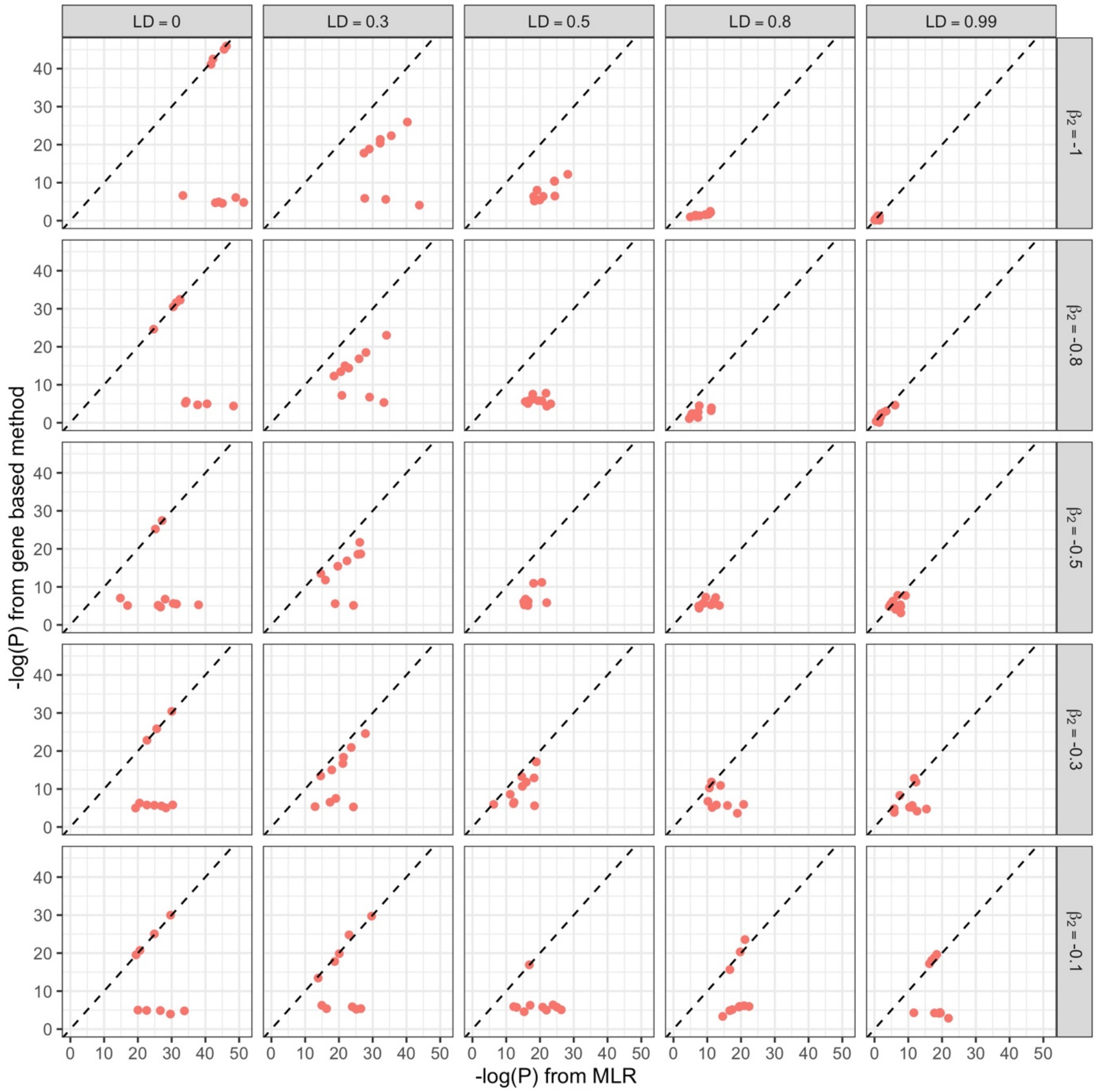


**Supplementary Figure 18.** Comparison of -log$_{10}$(*P*-value) from Imhof's method to compute the P value and use fastBAT as fallback with that from MLR as the gold standard in the scenario of masking effect. The results from fastBAT corresponding to the same scenario are shown in Supplementary Fig. 2.

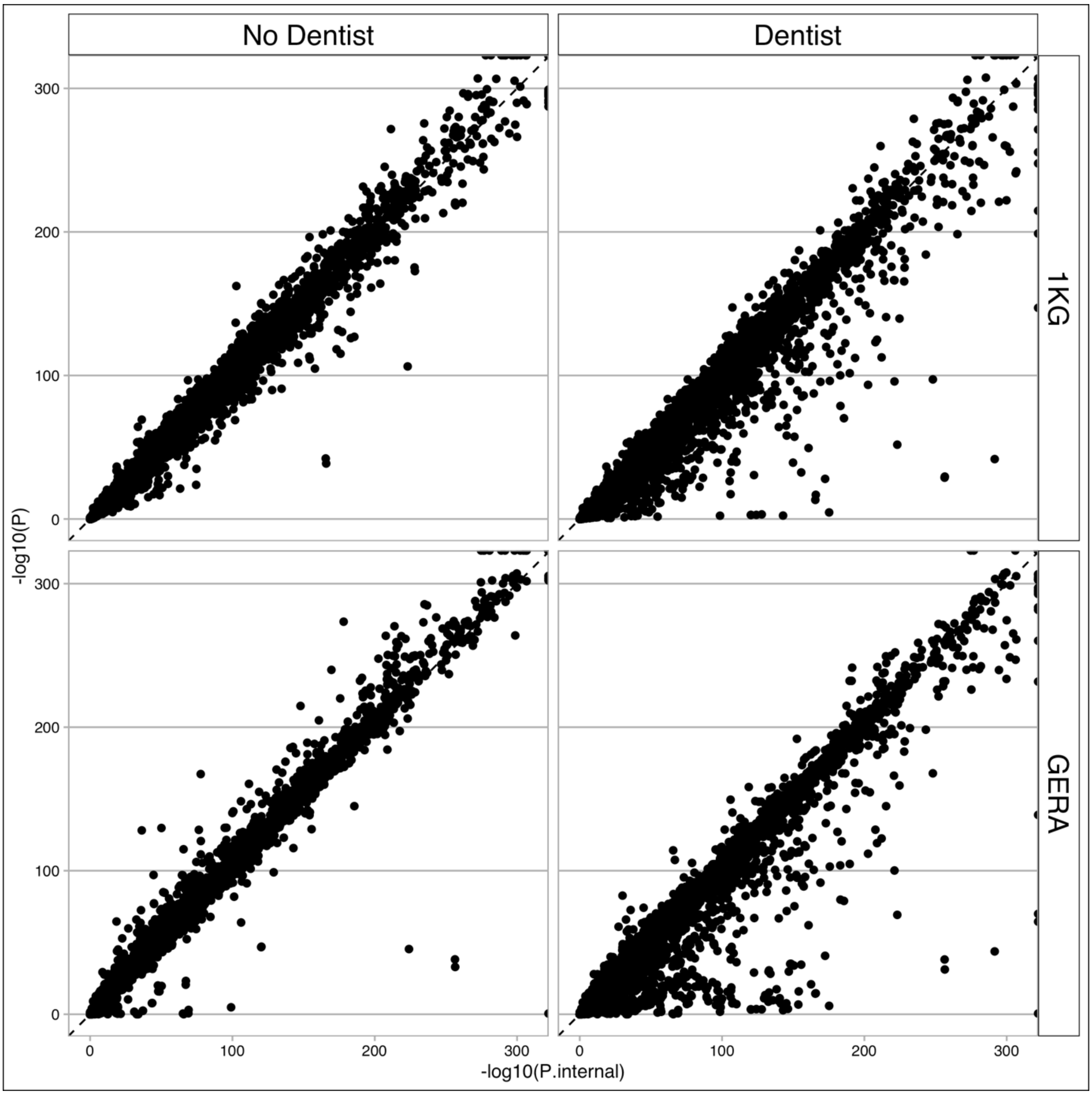


**Supplementary Figure 19.** Comparison of P-values of mBAT-combo between using in-sample LD (x-axis) and external LD (y-axis) in the simulations based on real genotype data. Rows are results using different LD reference data: GERA and 1KG phase3 of European ancestry. Columns are results without or with DENTIST application on the GWAS summary data to capture SNPs with inconsistent LD between GWAS and reference samples.

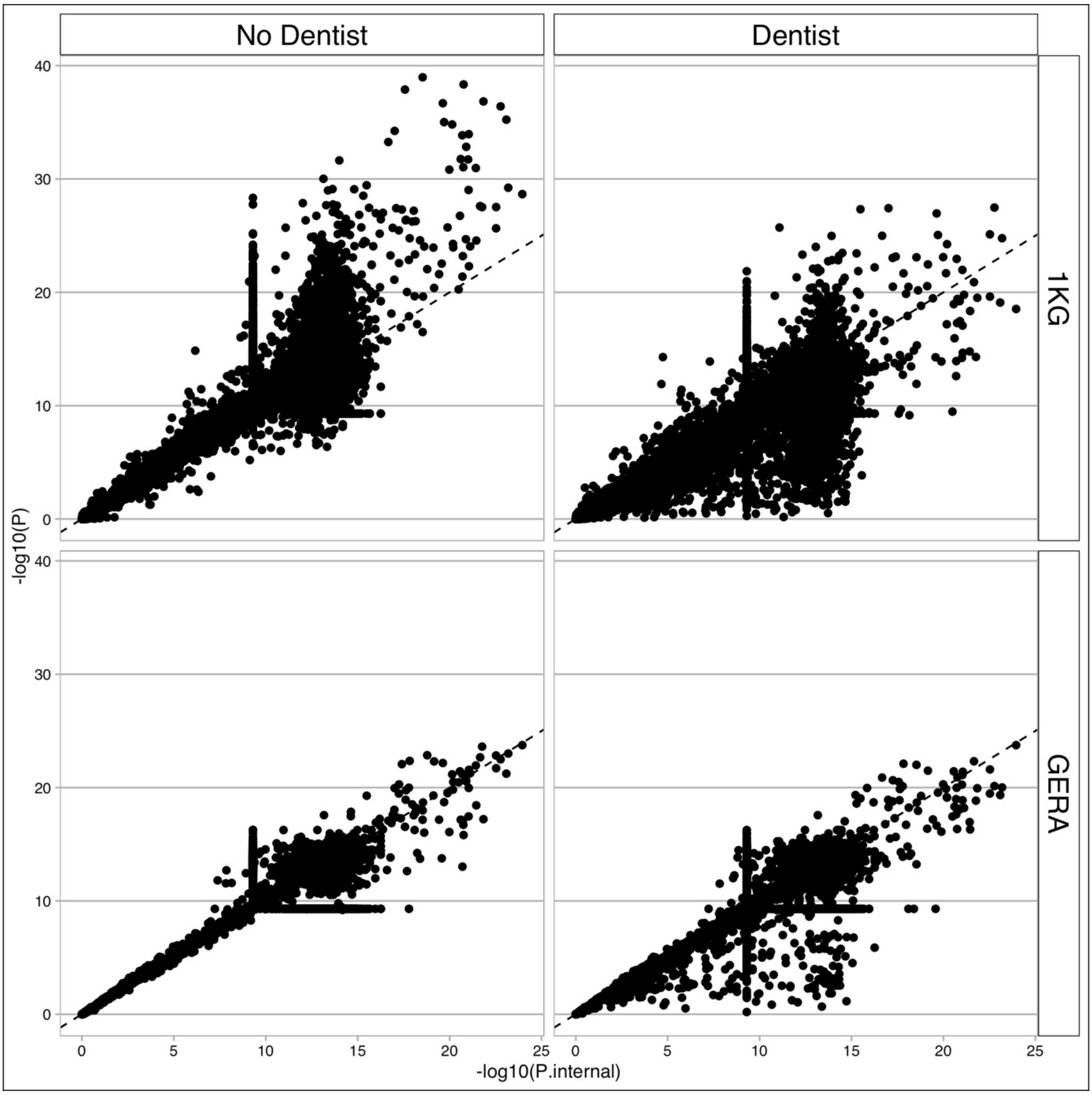


**Supplementary Figure 20** Comparison of P-values of MAGMA between using in-sample LD (x-axis) and external LD (y-axis) in the simulations based on real genotype data. Rows are results using different LD reference data: GERA and 1KG phase3 of European ancestry. Columns are results without or with DENTIST application on the GWAS summary data to capture SNPs with inconsistent LD between GWAS and reference samples. The horizontal and vertical lines reflect a result of the fallback strategy for the Imhof's method to compute the P-values in MAGMA.

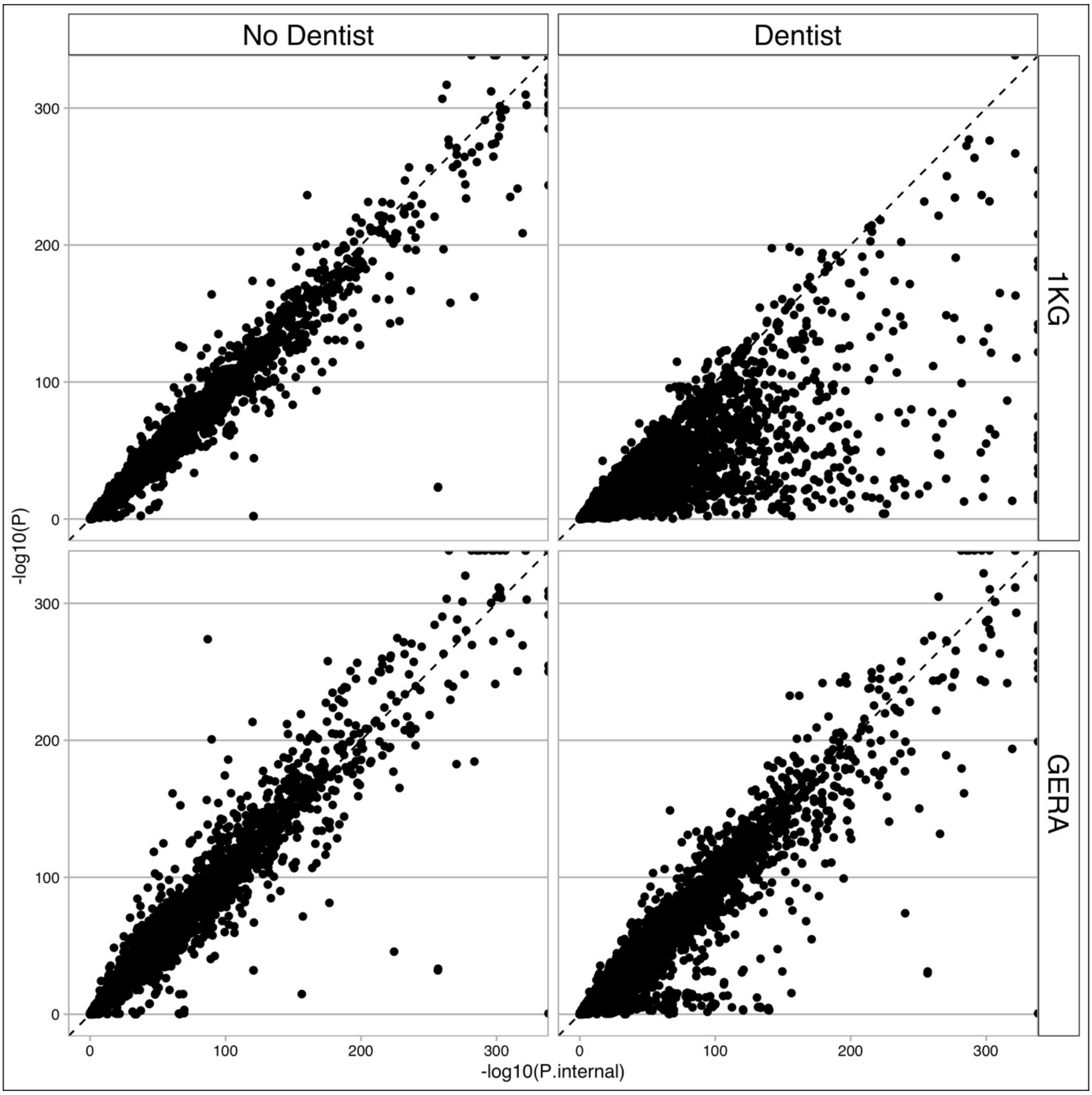


**Supplementary Figure 21** Comparison of P-values of fastBAT between using in-sample LD (x-axis) and external LD (y-axis) in the simulations based on real genotype data. Rows are results using different LD reference data: GERA and 1KG phase3 of European ancestry. Columns are results without or with DENTIST application on the GWAS summary data to capture SNPs with inconsistent LD between GWAS and reference samples.

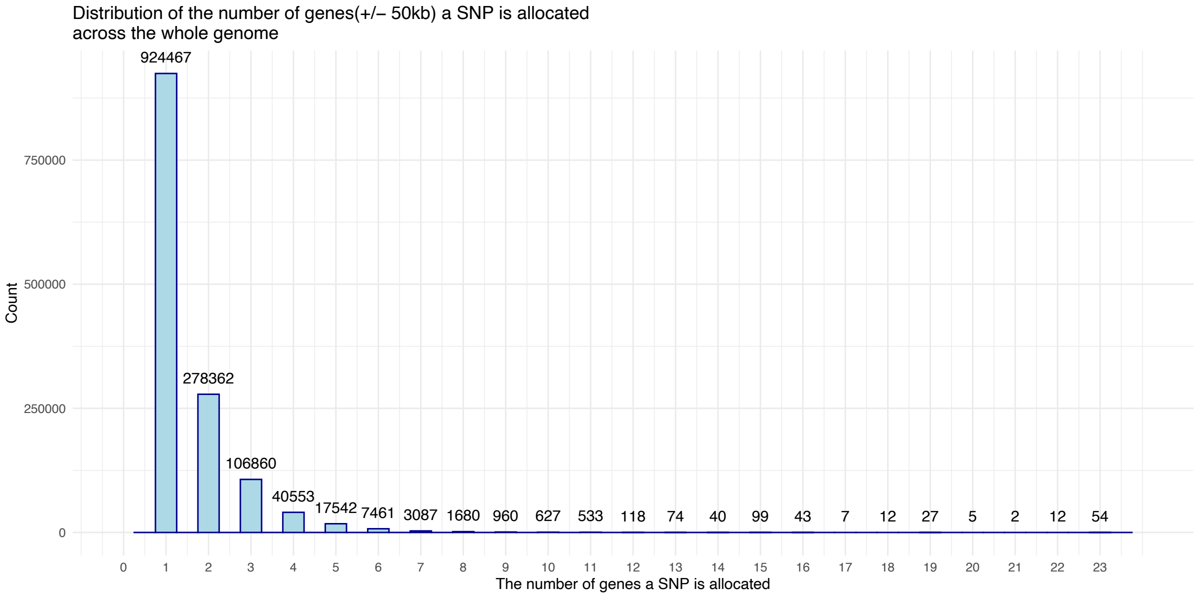


**Supplementary Figure 22** Distribution of the numbers of genes a SNP is allocated based on whether its physical position is within a gene region ± 50kb. Results are based on the GERA imputed SNPs (MAF > 0.01, European population).

| Trait | Gene | $P_{mBAT}$ | $P_{fastBAT}$ | $P_{MAGMA}$ | $\beta_1$ | $P_{\beta 1}$ | $\beta_2$ | $P_{\beta 2}$ | $\beta_1 \times \beta_2 \times$ LD |
|---|---|---|---|---|---|---|---|---|---|
| AST ALT ratio | *VSTM5* | 4.36E-26 | 2.17E-05 | 5.47E-05 | 0.0280891 | 3.47E-19 | 0.0193899 | 6.40E-10 | -0.0003518 |
| AST ALT ratio | *HLA-DQA2* | 1.89E-08 | 6.63E-06 | 0.00024683 | 0.0269455 | 2.15E-25 | 0.0395213 | 3.39E-24 | -0.0004017 |
| Albumin | *MARS* | 1.36E-12 | 2.37E-05 | 0.0072144 | 0.0199712 | 1.17E-12 | 0.0160355 | 1.25E-07 | -0.0001465 |
| Alkaline phosphatase | *LOC100996634* | 3.61E-11 | 1.45E-05 | 0.00018053 | 0.0242273 | 7.41E-13 | 0.0152519 | 5.48E-09 | -0.0001491 |
| Apolipoprotein A | *EIF2B1* | 7.57E-12 | 3.76E-06 | 0.00016202 | 0.0651307 | 4.10E-09 | 0.0119289 | 2.15E-06 | -9.40E-05 |
| Apolipoprotein A | *DHX37* | 4.98E-11 | 8.05E-05 | 0.045266 | 0.0751721 | 2.83E-34 | 0.0456573 | 1.35E-17 | -0.0027361 |
| Apolipoprotein A | *RAB3D* | 1.40E-08 | 2.27E-05 | 6.19E-06 | 0.0344501 | 3.75E-17 | 0.0418517 | 4.45E-07 | -0.0005597 |
| Apolipoprotein A | *FCRLA* | 3.44E-08 | 3.81E-06 | 0.0001162 | 0.0208618 | 6.58E-11 | 0.0166546 | 6.94E-09 | -0.00012 |
| Apolipoprotein A | *UBE2D3* | 4.23E-11 | 1.11E-05 | 0.03734 | 0.0177906 | 1.05E-09 | -0.040432 | 3.35E-09 | -7.78E-05 |
| Apolipoprotein A | *CISD2* | 4.63E-09 | 1.30E-05 | 0.051723 | 0.0177906 | 1.05E-09 | -0.040432 | 3.35E-09 | -7.78E-05 |
| Apolipoprotein A | *SLC9B1* | 5.58E-12 | 0.00012242 | 0.040486 | 0.0177906 | 1.05E-09 | -0.040432 | 3.35E-09 | -7.78E-05 |
| Apolipoprotein A | *SLC22A3* | 2.36E-12 | 3.79E-06 | 0.00022211 | 0.0479315 | 3.99E-15 | 0.0160518 | 1.16E-06 | -0.0001376 |
| Apolipoprotein A | *DEFB135* | 1.13E-10 | 3.28E-06 | 3.43E-06 | 0.184801 | 2.05E-127 | 0.108577 | 5.84E-64 | -0.0183954 |
| Apolipoprotein A | *ZNF696* | 1.39E-14 | 0.00130231 | 0.01284 | 0.0311966 | 2.71E-16 | 0.0308778 | 1.02E-06 | -7.35E-05 |
| Apolipoprotein A | *COL27A1* | 2.60E-10 | 5.89E-06 | 1.17E-05 | 0.0167055 | 8.42E-11 | 0.0279909 | 5.34E-08 | -0.000111 |
| Apolipoprotein A | *OBP2B* | 3.36E-10 | 0.0001154 | 0.0074248 | 0.0291542 | 4.45E-18 | 0.0435667 | 9.58E-17 | -0.0002896 |
| Apolipoprotein B adjstatins | *CYP4F2* | 1.33E-08 | 0.00018156 | 9.05E-05 | 0.0148209 | 1.51E-09 | 0.0127471 | 2.02E-07 | -3.94E-05 |
| Apolipoprotein B adjstatins | *PDLIM4* | 2.76E-15 | 1.04E-05 | 0.00026887 | 0.0174559 | 6.99E-11 | 0.0149514 | 2.33E-08 | -6.23E-05 |
| Apolipoprotein B adjstatins | *ZNRD1* | 1.40E-11 | 4.50E-06 | 9.96E-05 | 0.0197848 | 7.21E-16 | 0.0159694 | 7.45E-11 | -6.52E-05 |
| Apolipoprotein B adjstatins | *PPP1R11* | 1.48E-10 | 7.14E-06 | 0.0004314 | 0.0197848 | 7.21E-16 | 0.0159694 | 7.45E-11 | -6.52E-05 |
| Apolipoprotein B adjstatins | *RNF39* | 1.98E-09 | 1.48E-05 | 0.00063512 | 0.0197848 | 7.21E-16 | 0.0159694 | 7.45E-11 | -6.52E-05 |
| Apolipoprotein B adjstatins | *TRIM26* | 4.88E-11 | 8.77E-06 | 0.039884 | 0.0366863 | 1.76E-21 | -0.018832 | 2.31E-13 | -0.0002532 |
| Aspartate aminotransferase | *TMEM40* | 2.93E-11 | 7.95E-05 | 0.00011899 | 0.0183182 | 2.50E-12 | 0.0187327 | 1.61E-10 | -0.0001012 |
| Aspartate aminotransferase | *OR12D3* | 1.86E-09 | 0.00010323 | 0.0020356 | 0.0283375 | 1.49E-23 | -0.040018 | 4.83E-11 | -0.0003445 |
| Aspartate aminotransferase | *TRIM26* | 9.80E-13 | 8.00E-06 | 0.027256 | 0.0344575 | 2.39E-29 | 0.0354357 | 6.91E-25 | -0.0003796 |
| C reactive protein | *EGFL8* | 4.37E-10 | 3.41E-06 | 4.55E-05 | 0.0337403 | 1.69E-20 | -0.01683 | 5.44E-11 | -0.0002052 |
| C reactive protein | *AGPAT1* | 1.06E-09 | 4.66E-06 | 7.13E-05 | 0.0337403 | 1.69E-20 | -0.01683 | 5.44E-11 | -0.0002052 |
| C reactive protein | *OBP2B* | 9.02E-16 | 3.72E-05 | 0.00011715 | 0.0246796 | 4.63E-18 | 0.0262616 | 2.54E-16 | -0.0003509 |
| Calcium | *HLA-B* | 2.18E-09 | 0.00012112 | 3.02E-05 | 0.0137778 | 4.54E-08 | 0.0119096 | 2.28E-06 | -2.04E-05 |
| Calcium | *MICA* | 7.93E-11 | 0.00027062 | 0.00010767 | 0.0137778 | 4.54E-08 | 0.0119096 | 2.28E-06 | -2.04E-05 |
| Cholesterol adjstatins | *SLC29A2* | 1.36E-09 | 4.89E-05 | 0.096175 | 0.0158579 | 1.32E-10 | 0.0126947 | 2.69E-07 | -4.77E-05 |
| Cholesterol adjstatins | *MLEC* | 2.11E-15 | 6.00E-06 | 2.29E-05 | 0.0159786 | 3.90E-11 | 0.0126296 | 5.33E-07 | -2.51E-05 |
| Cholesterol adjstatins | *CYP4F2* | 8.56E-12 | 4.28E-05 | 1.17E-05 | 0.0152931 | 4.50E-10 | 0.0150442 | 8.55E-10 | -4.80E-05 |
| Cholesterol adjstatins | *RNF39* | 8.10E-13 | 3.77E-06 | 9.80E-05 | 0.0234286 | 4.39E-21 | 0.0208689 | 7.77E-16 | -0.0001298 |
| Cystatin C | *C6orf47* | 2.64E-09 | 3.51E-06 | 8.88E-05 | 0.0314552 | 1.71E-35 | 0.0144275 | 1.17E-08 | -0.0001441 |
| Gamma glutamyltransferase | *HLA-DRB5* | 1.38E-09 | 0.00050532 | 2.11E-05 | 0.0220155 | 1.28E-17 | 0.0198673 | 7.30E-09 | -0.0001623 |
| Glucose | *OBP2B* | 1.57E-14 | 1.12E-05 | 2.15E-05 | 0.0210287 | 2.89E-13 | 0.0217028 | 1.74E-11 | -0.0002288 |
| Glycated haemoglobin HbA1c | *MARS* | 3.49E-10 | 7.23E-06 | 2.22E-05 | 0.0199453 | 2.65E-13 | 0.0141707 | 1.83E-08 | -8.68E-05 |
| Glycated haemoglobin HbA1c | *CDK5RAP1* | 2.27E-08 | 9.10E-06 | 0.00057641 | 0.0275161 | 9.60E-11 | -0.013685 | 8.06E-08 | -0.0001315 |
| Glycated haemoglobin HbA1c | *SLC22A5* | 9.28E-09 | 0.00024418 | 0.0096619 | -0.03179 | 5.11E-09 | 0.0141647 | 1.24E-06 | -5.61E-05 |
| Glycated haemoglobin HbA1c | *HLA-DRA* | 2.94E-11 | 0.00029974 | 0.0010938 | 0.0364901 | 1.96E-27 | 0.0216329 | 5.38E-15 | -0.0003609 |
| IGF 1 | *UQCC1* | 3.51E-11 | 0.00129586 | 0.0043006 | 0.0281008 | 1.93E-10 | 0.0116362 | 2.32E-06 | -7.56E-05 |
| IGF 1 | *MUC21* | 8.18E-10 | 2.45E-05 | 0.00024201 | -0.015293 | 9.88E-10 | 0.023454 | 3.23E-09 | -0.0001016 |
| LDL direct adjstatins | *CYP4F2* | 5.08E-11 | 6.13E-05 | 2.31E-05 | 0.0152287 | 5.32E-10 | 0.014731 | 1.89E-09 | -4.68E-05 |
| LDL direct adjstatins | *PDLIM4* | 2.08E-13 | 8.29E-05 | 0.0034943 | 0.0180672 | 1.49E-11 | 0.0128967 | 1.45E-06 | -5.56E-05 |
| LDL direct adjstatins | *ZNRD1* | 1.23E-14 | 3.57E-06 | 6.83E-05 | 0.02124 | 4.71E-18 | 0.0186687 | 2.70E-14 | -8.18E-05 |
| LDL direct adjstatins | *PPP1R11* | 1.50E-13 | 4.69E-06 | 0.00043828 | 0.02124 | 4.71E-18 | 0.0186687 | 2.70E-14 | -8.18E-05 |
| LDL direct adjstatins | *RNF39* | 8.13E-12 | 7.94E-06 | 0.00057205 | 0.02124 | 4.71E-18 | 0.0186687 | 2.70E-14 | -8.18E-05 |
| Non albumin protein | *LOC645177* | 4.85E-08 | 1.23E-05 | 4.22E-06 | 0.0226406 | 7.43E-09 | 0.0127714 | 1.00E-06 | -2.69E-05 |
| Non albumin protein | *LRMP* | 3.38E-08 | 6.19E-05 | 0.00017818 | 0.0226406 | 7.43E-09 | 0.0127714 | 1.00E-06 | -2.69E-05 |
| Non albumin protein | *HIST1H4E* | 5.54E-09 | 1.01E-05 | 0.0095073 | 0.0342725 | 1.54E-18 | 0.0294062 | 4.73E-14 | -0.0007872 |
| Non albumin protein | *OR2H2* | 8.85E-09 | 4.14E-06 | 2.85E-06 | 0.0398334 | 1.11E-34 | 0.0381882 | 1.74E-34 | -0.0009221 |
| Non albumin protein | *TRIM10* | 1.19E-08 | 6.75E-06 | 0.0042286 | 0.0308148 | 7.15E-32 | 0.0221897 | 3.39E-18 | -0.0001011 |
| Non albumin protein | *OBP2B* | 1.29E-08 | 0.00419226 | 0.018626 | 0.0227611 | 4.42E-15 | 0.0200091 | 2.79E-09 | -0.0002326 |
| SHBG | *IFI27* | 9.50E-14 | 1.31E-05 | 3.97E-05 | 0.0508046 | 3.49E-30 | -0.043129 | 3.27E-22 | -0.0018112 |
| SHBG | *CATIP* | 1.70E-08 | 5.97E-05 | 2.45E-05 | 0.0143129 | 1.29E-08 | 0.0124273 | 7.87E-07 | -2.05E-05 |
| Testosterone | *DCAF12* | 2.93E-10 | 1.10E-05 | 0.00023783 | 0.0181311 | 2.89E-10 | 0.0161749 | 1.82E-06 | -6.70E-05 |
| Total bilirubin | *EHMT2* | 5.51E-15 | 4.76E-06 | 1.18E-05 | 0.0399194 | 1.08E-24 | 0.0143991 | 1.58E-07 | -0.0002798 |
| Total bilirubin | *C2* | 2.46E-12 | 6.23E-06 | 2.47E-05 | 0.0399193 | 1.08E-24 | 0.0143991 | 1.58E-07 | -0.0002798 |
| Total bilirubin | *ZBTB12* | 2.35E-14 | 1.19E-05 | 1.42E-05 | 0.0399193 | 1.08E-24 | 0.0143991 | 1.58E-07 | -0.0002798 |
| Total protein | *SNX29* | 6.79E-10 | 0.00100438 | 0.00084043 | 0.0141338 | 1.83E-08 | 0.0234277 | 1.94E-06 | -3.36E-05 |
| Total protein | *OSM* | 9.46E-09 | 0.00125967 | 0.00023936 | 0.0257462 | 2.68E-09 | 0.0117619 | 2.93E-06 | -3.41E-05 |
| Total protein | *TMEM40* | 4.95E-08 | 0.0149277 | 0.0057771 | 0.0193368 | 1.99E-10 | -0.013589 | 6.14E-07 | -7.91E-05 |
| Urate | *AGAP11* | 4.99E-17 | 1.94E-05 | 2.82E-06 | 0.0313151 | 4.43E-11 | 0.0133823 | 7.09E-08 | -9.50E-05 |
| Urate | *LOC101927789* | 8.84E-13 | 1.12E-05 | 0.0027591 | 0.0335129 | 1.11E-13 | 0.0428773 | 1.10E-11 | -7.10E-05 |
| Urate | *FUK* | 4.00E-17 | 3.49E-06 | 0.0026307 | 0.0228567 | 7.02E-10 | 0.0377978 | 3.43E-07 | -5.30E-05 |
| Urate | *COG4* | 1.01E-19 | 1.69E-05 | 0.024826 | 0.0216014 | 1.34E-08 | 0.0392836 | 3.43E-07 | -3.24E-05 |
| Urate | *TCF4* | 2.64E-08 | 8.22E-06 | 5.37E-05 | 0.0356241 | 4.34E-11 | 0.0385734 | 3.94E-07 | -5.26E-05 |
| Urate | *NBAS* | 4.07E-17 | 2.89E-05 | 4.94E-05 | 0.0237159 | 7.51E-16 | 0.0242368 | 8.97E-10 | -9.75E-05 |
| Urate | *ALS2CL* | 3.31E-08 | 0.0001484 | 0.0065986 | 0.0583167 | 3.83E-08 | 0.0220068 | 1.74E-06 | -4.12E-05 |
| Urate | *TMIE* | 4.77E-10 | 0.00029908 | 0.016265 | 0.0583167 | 3.83E-08 | 0.0220068 | 1.74E-06 | -4.12E-05 |
| Urate | *PRSS50* | 4.12E-10 | 0.00033851 | 0.015975 | 0.0583167 | 3.83E-08 | 0.0220068 | 1.74E-06 | -4.12E-05 |
| Urate | *PRSS46* | 2.02E-10 | 0.00019236 | 0.018262 | 0.0583167 | 3.83E-08 | 0.0220068 | 1.74E-06 | -4.12E-05 |
| Urate | *PRSS45* | 3.41E-10 | 0.00085211 | 0.044881 | 0.0583167 | 3.83E-08 | 0.0220068 | 1.74E-06 | -4.12E-05 |
| Urate | *CCDC51* | 1.09E-12 | 2.77E-06 | 6.87E-05 | 0.0168839 | 1.92E-08 | 0.0578798 | 6.62E-08 | -5.42E-05 |
| Urate | *TMA7* | 2.16E-14 | 2.96E-06 | 6.17E-05 | 0.0168839 | 1.92E-08 | 0.0578798 | 6.62E-08 | -5.42E-05 |
| Urate | *ATRIP* | 5.99E-16 | 4.70E-06 | 4.03E-05 | 0.0168839 | 1.92E-08 | 0.0578798 | 6.62E-08 | -5.42E-05 |
| Urate | *TREX1* | 1.10E-16 | 1.05E-05 | 3.39E-05 | 0.0168839 | 1.92E-08 | 0.0578798 | 6.62E-08 | -5.42E-05 |
| Urate | *PFKFB4* | 7.58E-19 | 5.32E-06 | 0.0030428 | 0.0307076 | 6.81E-16 | 0.0530711 | 8.49E-08 | -6.57E-05 |
| Urate | *CELSR3* | 2.89E-08 | 0.0001844 | 0.023864 | 0.0238118 | 2.63E-12 | 0.0540555 | 8.97E-08 | -5.74E-05 |
| Urate | *CACNA2D2* | 7.06E-17 | 1.93E-05 | 0.037928 | 0.037509 | 8.04E-17 | 0.0556522 | 2.60E-07 | -6.43E-05 |
| Urate | *GRM2* | 4.16E-17 | 1.48E-05 | 0.010607 | 0.0396712 | 2.07E-18 | 0.0191675 | 1.25E-07 | -8.58E-05 |
| Urate | *ODF1* | 3.75E-10 | 0.00037817 | 0.00048882 | 0.0163384 | 1.11E-10 | 0.0145194 | 9.89E-09 | -7.68E-05 |
| Urea | *CAT* | 2.61E-09 | 0.0001053 | 0.00019212 | 0.0197279 | 1.39E-14 | 0.0204898 | 4.20E-08 | -0.0001439 |
| Urea | *ST3GAL2* | 6.64E-17 | 3.20E-06 | 0.0003389 | 0.0262955 | 1.29E-08 | 0.0183304 | 8.27E-07 | -4.88E-05 |
| Urea | *FUK* | 3.67E-15 | 2.08E-05 | 0.007509 | 0.0262836 | 1.31E-08 | 0.0181406 | 1.07E-06 | -4.83E-05 |
| Urea | *COG4* | 3.47E-17 | 3.77E-05 | 0.043714 | 0.0262403 | 1.38E-08 | 0.0179221 | 1.44E-06 | -4.75E-05 |
| Urea | *ATP9B* | 6.56E-10 | 2.81E-06 | 7.62E-05 | 0.0252609 | 1.17E-10 | 0.0184203 | 7.31E-07 | -4.91E-05 |
| Urea | *MISP* | 1.33E-08 | 3.18E-06 | 2.92E-05 | -0.020962 | 1.79E-08 | 0.0169688 | 1.44E-06 | -3.93E-05 |
| Urea | *GPR4* | 2.72E-14 | 4.00E-05 | 0.0010325 | 0.0534169 | 2.06E-19 | 0.0275346 | 1.22E-08 | -7.84E-05 |
| Urea | *NBAS* | 4.69E-08 | 0.0107494 | 0.12939 | 0.0225659 | 2.29E-11 | 0.0166507 | 1.97E-08 | -8.05E-05 |
| Urea | *ARGFX* | 2.51E-08 | 1.85E-05 | 0.00089272 | 0.0217213 | 6.51E-15 | 0.0150205 | 5.59E-07 | -0.0001191 |
| Urea | *SLC34A1* | 1.43E-17 | 8.09E-06 | 0.0011286 | 0.0266241 | 9.23E-11 | 0.0331921 | 7.61E-07 | -4.84E-05 |
| Urea | *PFN3* | 1.59E-16 | 4.42E-05 | 0.0021398 | 0.0256334 | 4.17E-10 | 0.030638 | 1.17E-06 | -4.54E-06 |
| Urea | *F12* | 8.77E-17 | 2.88E-05 | 0.0015473 | 0.0256334 | 4.17E-10 | 0.030638 | 1.17E-06 | -4.54E-06 |

| | | | | | | | | | |
|---|---|---|---|---|---|---|---|---|---|
| Urea | *GRK6* | 5.70E-16 | 5.04E-06 | 0.00050403 | 0.0256334 | 4.17E-10 | 0.030638 | 1.17E-06 | -4.54E-06 |
| Vitamin D | *UTP3* | 1.14E-14 | 2.80E-05 | 0.001156 | 0.0190853 | 2.53E-14 | 0.0139499 | 3.45E-06 | -1.63E-05 |
| eGFR | *EFNA1* | 3.32E-12 | 7.76E-06 | 5.25E-05 | 0.0250579 | 2.01E-10 | 0.0125797 | 8.53E-07 | -0.0001087 |
| AST ALT ratio | *VSTM5* | 4.36E-26 | 2.17E-05 | 5.47E-05 | 0.0280891 | 3.47E-19 | 0.0193899 | 6.40E-10 | -0.0003518 |
| AST ALT ratio | *HLA-DQA2* | 1.89E-08 | 6.63E-06 | 0.00024683 | 0.0269455 | 2.15E-25 | 0.0395213 | 3.39E-24 | -0.0004017 |
| Albumin | *MARS* | 1.36E-12 | 2.37E-05 | 0.0072144 | 0.0199712 | 1.17E-12 | 0.0160355 | 1.25E-07 | -0.0001465 |
| Alkaline phosphatase | *LOC100996634* | 3.61E-11 | 1.45E-05 | 0.00018053 | 0.0242273 | 7.41E-13 | 0.0152519 | 5.48E-09 | -0.0001491 |
| Apolipoprotein A | *EIF2B1* | 7.57E-12 | 3.76E-06 | 0.00016202 | 0.0651307 | 4.10E-09 | 0.0119289 | 2.15E-06 | -9.40E-05 |
| Apolipoprotein A | *DHX37* | 4.98E-11 | 8.05E-05 | 0.045266 | 0.0751721 | 2.83E-34 | 0.0456573 | 1.35E-17 | -0.0027361 |
| Apolipoprotein A | *RAB3D* | 1.40E-08 | 2.27E-05 | 6.19E-06 | 0.0344501 | 3.75E-17 | 0.0418517 | 4.45E-07 | -0.0005597 |
| Apolipoprotein A | *FCRLA* | 3.44E-08 | 3.81E-06 | 0.0001162 | 0.0208618 | 6.58E-11 | 0.0166546 | 6.94E-09 | -0.00012 |
| Apolipoprotein A | *UBE2D3* | 4.23E-11 | 1.11E-05 | 0.03734 | 0.0177906 | 1.05E-09 | -0.040432 | 3.35E-09 | -7.78E-05 |
| Apolipoprotein A | *CISD2* | 4.63E-09 | 1.30E-05 | 0.051723 | 0.0177906 | 1.05E-09 | -0.040432 | 3.35E-09 | -7.78E-05 |
| Apolipoprotein A | *SLC9B1* | 5.58E-12 | 0.00012242 | 0.040486 | 0.0177906 | 1.05E-09 | -0.040432 | 3.35E-09 | -7.78E-05 |
| Apolipoprotein A | *SLC22A3* | 2.36E-12 | 3.79E-06 | 0.00022211 | 0.0479315 | 3.99E-15 | 0.0160518 | 1.16E-06 | -1.71E-05 |
| Apolipoprotein A | *DEFB135* | 1.13E-10 | 3.28E-06 | 3.43E-06 | 0.184801 | 2.05E-127 | 0.108577 | 5.84E-64 | -0.0183954 |
| Apolipoprotein A | *ZNF696* | 1.39E-14 | 0.00130231 | 0.01284 | 0.0311966 | 2.71E-16 | 0.0308778 | 1.02E-06 | -7.35E-05 |
| Apolipoprotein A | *COL27A1* | 2.60E-10 | 5.89E-06 | 1.17E-05 | 0.0167055 | 8.42E-11 | 0.0279909 | 5.34E-08 | -0.000111 |
| Apolipoprotein A | *OBP2B* | 3.36E-10 | 0.0001154 | 0.0074248 | 0.0291542 | 4.45E-18 | 0.0435667 | 9.58E-17 | -0.0002896 |
| Apolipoprotein B adjstatins | *CYP4F2* | 1.33E-08 | 0.00018156 | 9.05E-05 | 0.0148209 | 1.51E-09 | 0.0127471 | 2.02E-07 | -3.94E-05 |
| Apolipoprotein B adjstatins | *PDLIM4* | 2.76E-15 | 1.04E-05 | 0.00026887 | 0.0174559 | 6.99E-11 | 0.0149514 | 2.33E-08 | -6.23E-05 |
| Apolipoprotein B adjstatins | *ZNRD1* | 1.40E-11 | 4.50E-06 | 9.96E-05 | 0.0197848 | 7.21E-16 | 0.0159694 | 7.45E-11 | -6.52E-05 |
| Apolipoprotein B adjstatins | *PPP1R11* | 1.48E-10 | 7.14E-06 | 0.0004314 | 0.0197848 | 7.21E-16 | 0.0159694 | 7.45E-11 | -6.52E-05 |

**Supplementary table 1**. Genes that were conservatively significant in mBAT ($P_{mBAT} < 5\times10^{-8}$) but missed by the other gene-based chi-square sum methods (both $P_{fastBAT}$ and $P_{MAGMA} > 2.7\times10^{-6}$), with masking effect associations detected by COJO in 23 UK Biobank blood and urine metabolites. $\beta_1$ and $\beta_2$ are the estimated joint effect sizes of the top 2 SNPs and $P_{\beta_1}$ and $P_{\beta_2}$ are the joint association P-values

| | COJO significant | COJO insignificant |
|---|---|---|
| **mBAT ($P<5\times10^{-8}$)** | 98 | 4,175 |
| **random genes** | 4,990 | 636,035 |

**Supplementary table 2**. Fisher's exact test for whether mBAT significant gene-trait pairs at a stringent significance level ($5\times10^{-8}$) was enriched in those with significant COJO signals with masking effect. The number of mBAT significant masking gene-trait pairs confirmed by COJO was 98. The number of mBAT significant pairs that were not detected by COJO was 4,175. The total number of gene-trait pairs was 641,025 = 18,315 protein coding genes $\times$ 35 traits, among which 4,990 pairs were significant in COJO with masking but not in mBAT stringent.
